# A Genome-Scale Virtual *Escherichia coli* in the L++ Language

**DOI:** 10.64898/2026.11.25.769137

**Authors:** Donghoon M. Lee, Hyun-Ju Cho, Kyong-Cheol Ko, Daehwan Kim

## Abstract

A virtual organism is a computational model, mechanistic or learned from data, that predicts how a living system grows, adapts and responds to genetic edits, drugs and environments. As a first step toward such a model, we are building a virtual *Escherichia coli*, and here we present its first version, *E. coli* version 1.0, written for K-12 MG1655 in the L++ programming language. The model writes the cell as a biologist reads it, in one cell definition that compiles to 33,246 species and 41,649 reactions over 4,484 gene loci. Every gene’s transcription, translation and decay are written out, for 4,196 of the 4,374 transcribed genes with the gene’s own nucleotide and residue counts from the genome sequence, so the proteins that give *E. coli* its identity and functions, from metabolic enzymes to the machinery that replicates, divides and builds the cell, are each made from their own gene, with the promoter constants of 2,016 genes set toward measured protein abundances. The code wires 360 of 377 essential-gene candidates into processes that growth needs, among them all 293 that at least two genome-wide screens call essential; 177 of the 360 are read by gates and slowdowns written from those screens’ calls, so for them the model’s requirement follows the screens by construction. Twenty-eight two-component systems are written, and on development versions the two built from measured constants, for osmolarity and phosphate, reproduce the direction of their responses and the promoter order their constants were set to give. In a modified rich medium the cell divides every 1,100 to 1,400 s, at a pace set by a replication rate chosen to fit a 20-minute cycle. Of 95 metabolite pools measured in glucose minimal medium, 78 start within tenfold of measurement, 82 lie within it over *t* = 1,000 to 3,000 s and 65 over *t* = 10,000 to 20,000 s, and the model states where it does not yet match, among them 31 pools held at fixed values. Because the organism is code, a deletion, a retuned promoter or an added module is an edit, so one model serves both to understand the cell and to design its variants, and we hope to extend it toward a complete virtual *E. coli* together with scientists who bring their expertise and measurements.

## Introduction

*Escherichia coli* is the reference organism of molecular biology. Its genome has been closed and annotated for more than a quarter of a century [1]; its metabolism has been reconstructed at genome scale and repeatedly refined [2, 3, 4, 5], and the steps of such a reconstruction are set out as a protocol [6]; its transcriptional regulation is curated in RegulonDB [7] and EcoCyc [8]; its pathways, enzymes and reaction kinetics are cataloged in KEGG [9], BRENDA [10], SABIO-RK [11], UniProt [12] and BiGG [13]; and its proteome and metabolome have been measured in absolute units [14, 15, 16]. This knowledge lives in incompatible representations (stoichiometric matrices, regulatory tables, kinetic constants, microscopy and prose), and assembling any part of it into a running, mechanistic model of the cell still takes substantial bespoke engineering.

Whole-cell models closed part of this gap. The *Mycoplasma genitalium* whole-cell model predicted phenotype from genotype [17]; a large-scale mechanistic *E. coli* model, which simulates the transcription of about 4,350 genes and represents the functions of 1,214 of them, cross-evaluated heterogeneous datasets [18]; and a kinetic model of the JCVI-syn3A minimal cell reproduced its observed doubling behavior [19]. In each, the process logic and the rate laws are written in general-purpose code, and the parameters and the lists of genes, reactions and complexes are kept in separate knowledge bases or files. Constraint-based methods [20, 21] scale to the whole network and describe steady-state flux capacity; regulation, dynamics and gene expression enter them through added constraints and extensions, such as regulatory flux balance analysis [22], dynamic flux balance analysis [23] and models of metabolism and expression [24].

Here the organism itself is the artifact. L++ is a programming language whose nouns are biological: molecules, reactions, compartments, cells and organisms are first-class constructs, so a model reads like biology and composes by containment [25]. We describe a genome-scale virtual *E. coli* written entirely in L++: one mechanistic, human-readable codebase that expresses every gene, carries cited kinetic constants for 126 of the 6,186 reactions outside the per-gene expression rows, and simulates growth, gene expression, metabolism and the response to gene deletions and to changes in the medium.

The model is written from experimental results and known mechanisms, but some of its parts are inferred where measurements are missing, and some of its values may be inaccurate: many rate constants are model values, and some pools are held at measured values in place of their synthesis. The code shown here is therefore incomplete. We mark where the model still departs from measurement, and each result should be read with the measurement and the version it rests on. Much of the agreement with measurement reported here was built in, through calibration, starting values, held pools or rules written from the data, and we mark each case beside the comparison it affects. Our most important goal is to complete this model together with other scientists and to validate it further by experiment.

## Results

### Writing an organism in L++

We summarize only the constructs the model relies on; the language, its compiler lppc and its simulator lpps are described in the companion paper [25]. The unit of a program is the *object*: a species, reaction, compartment or cell, written as a name and an optional brace block, and nested to mirror the containment hierarchy of biology. Integer quantities denote countable molecules and decimals denote concentrations (molar by default). The distinction matters in a bacterium, where a gene is present in one or two copies while metabolites sit at millimolar.

L++ is protein-centric: the DNA, transcript and folded product of a gene share one name through the dot attribute, so GapA.gene, GapA.mRNA and GapA are the gene, its mRNA (both integer counts) and the enzyme. Because expression and catalysis refer to the same object, one declaration serves both roles (Figure 1). A small set of operators carries the structural meaning (Table 1): the tilde ∼ for a *has-a* complex such as RNAP∼RpoD, the colon : for an *is-a* state subtype such as protein:misfolded, the loaded pool [:] for a reservoir held constant, and genome positions written genome{n}.

**Figure 1.**
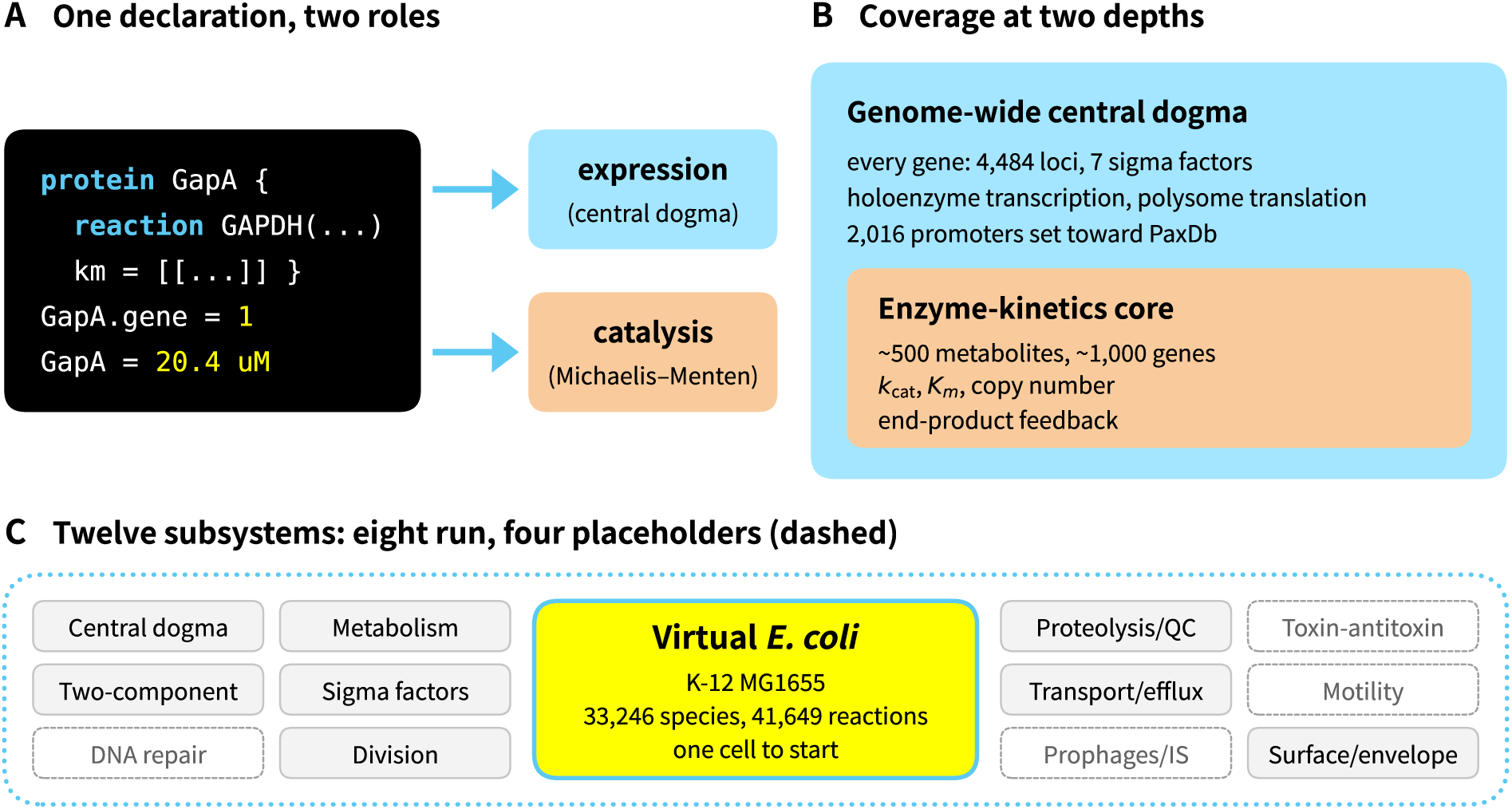
Architecture of the virtual *E. coli*. (A) Because expression and catalysis refer to one protein-centric object, a single declaration (here GapA) serves both its transcription and its enzymatic rate law. (B) Coverage at two depths, a genome-wide central-dogma layer over every gene and, within it, a kinetic core of about 500 metabolites and 1,000 genes carrying *k*_cat_, *K_m_* and copy number; of the 6,186 reactions outside the per-gene steps, 126 cite a source for some or all of their values (Methods). In version 1.0 the *σ* ^70^ holoenzyme transcribes nearly every gene, and the other six sigma factors transcribe eight genes in all. (C) The twelve subsystems of Table 2 compose by containment into one cell. Version 1.0 runs eight of them; DNA repair, toxin–antitoxin, motility and the prophages are present as expressed genes with placeholder or switched-off reactions.

**Table 1.** L++ operators used throughout the model.

| Token | Reads as | Example |
|---|---|---|
| -> | reactants yield products | A + B -> C |
| ~ | has-a: a bound complex | RNAP~RpoD |
| : | is-a: a state subtype | protein:misfolded |
| . | attribute of a species | GapA.gene |
| [:] | external or loaded pool | glucose[:] = 3.5 mM |
| .* | flat-merge import | Glycolysis.* |
| genome{n} | a genome position; {n:lo:hi} expands a reaction over positions lo to hi | genome{n} |

**Table 2.** Subsystems of the virtual *E. coli* (K-12 MG1655) and what version 1.0 runs of each. Every gene is expressed through the genome-wide central-dogma layer; the first eight subsystems also run their own reactions, and the last four are present as expressed genes with placeholder or switched-off reactions.

| Subsystem | What version 1.0 runs |
|---|---|
| *Run in version 1.0* |  |
| Central dogma | Holoenzyme transcription, polysome translation, tRNA charging, ribosome assembly; every gene expressed, with the promoter constants of 2,016 genes set toward their PaxDb levels |
| Metabolism | Central carbon, amino acids, cofactors, lipids and nucleotides as Michaelis–Menten enzymes (about 500 metabolites), with end-product feedback. The pentose phosphate pathway is written but switched off; glucose enters by an ATP-driven route and by phosphotransfer, with the glucose phosphotransferase system written in part |
| Two-component systems | 28 sensor kinase and response regulator pairs: EnvZ/OmpR and PhoR/PhoB from measured constants, 24 from one template and 2 on an older template |
| Sigma factors | Seven sigma factors bind one core RNA polymerase pool; *σ*<sup>70</sup> transcribes nearly every gene, and the alternative sigmas transcribe eight genes in all |
| Division and replication | Replication that needs the replisome, one round at a time from an origin held loaded; division at 4 *µ*m that needs the divisome. No DnaA-ATP control of initiation, and the Min assembly rows run at zero rate |
| Proteolysis and quality control | AAA+ proteases and chaperone triage of misfolded protein |
| Transport and efflux | Transporters for the nutrients and ions of the medium; the efflux pumps are expressed, and no drug efflux is evaluated |
| Surface and envelope | Cell wall built by the peptidoglycan pathway on measured kinetics; envelope synthesis that needs its essential enzymes; outer and inner membrane systems |
| *Placeholders in version 1.0* |  |
| DNA damage and repair | Repair and nucleoid genes expressed; the SOS module is written but switched off, and no repair or supercoiling reaction runs |
| Toxin–antitoxin | MazE–MazF and RelB–RelE binding that neutralizes each toxin; no reaction gives a toxin its activity |
| Motility and chemotaxis | CheA/CheY on the older template, with the sign of its signal and its output written in error (Results); flagellar assembly rows at zero rate and no flagellar rotation |
| Prophages and mobile elements | Genes expressed; the prophage modules are written but switched off, and no transposition reaction runs |

Reactions set their kinetics by named parameters, and each maps to a closed-form rate law. A catalyzing protein *E* multiplies the whole rate and defaults to 1 when omitted. With k and kr a reaction is reversible mass action,

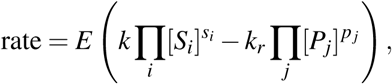

and adding km (with optional ks, ki and Hill exponent *n*) promotes it to a general enzyme law that collapses, for a single substrate, to Michaelis–Menten kinetics, rate = *Ek*[*S*]*/*(*K_m_* + [*S*]) with *k* read as *k*_cat_. In this law only the species that the km list names enter the rate; the other reactants and products are consumed and made at that rate without changing it (Methods). A count flux vc, optionally computed by a vc_function, drives directed processes such as polymerization. Metabolic enzymes are written as protein blocks carrying *k*_cat_, *K_m_* and, where relevant, product inhibition. The compiler checks syntax, names and units; it does not check elemental or charge balance, so each equation’s balance is kept by its author, and during a run lpps counts and reports any amount driven below zero, after redrawing or scaling the steps that would cause one (Methods).

### Model architecture

The virtual *E. coli* is grounded on the K-12 MG1655 reference genome [1, 26] (GenBank U00096.3). One cell definition, EcoliWholeCell, imports the modules of twelve subsystems, of which version 1.0 runs eight and holds four as placeholders (Table 2), and compiles to 33,246 species and 41,649 reactions over 4,484 gene loci and 4,377 transcripts; 35,463 of the reactions are the per-gene steps of the central dogma. The cell is a rod born at 2 *µ*m that divides at 4 *µ*m, with a periplasm around its cytoplasm, and lpps advances it in ticks of 2*^−^*^6^ s. Coverage runs at two depths (Figure 1): a genome-wide layer over every gene, and within it a core with enzyme kinetics. The cell imports each module by its place in the model library, and where two modules set the same value the later import wins: the expression layer that sets promoter constants toward PaxDb levels comes after the modules that type or tune a gene, the translation factors come after it, and the four files that write out each gene’s central dogma come last (Listing 1).

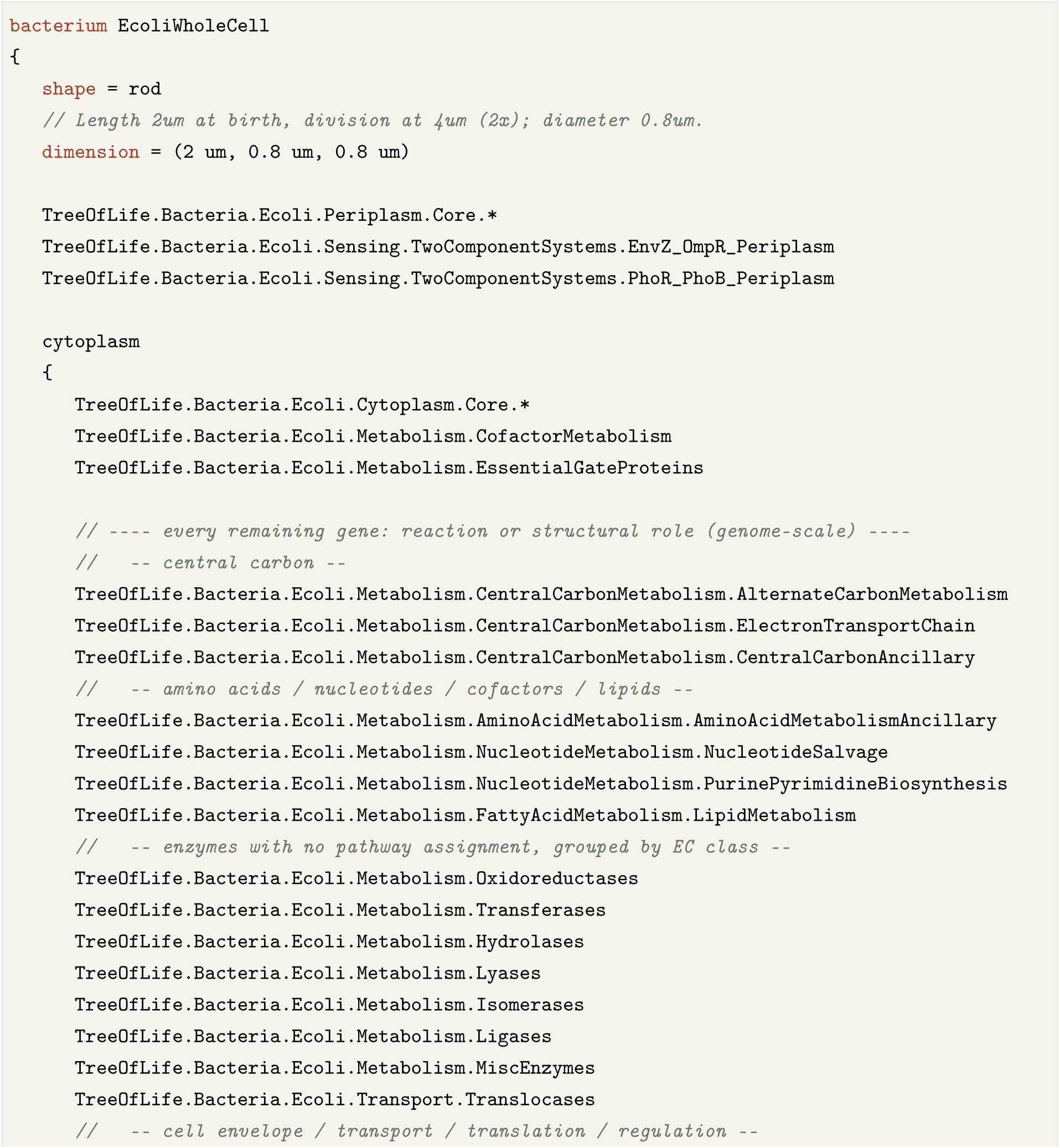
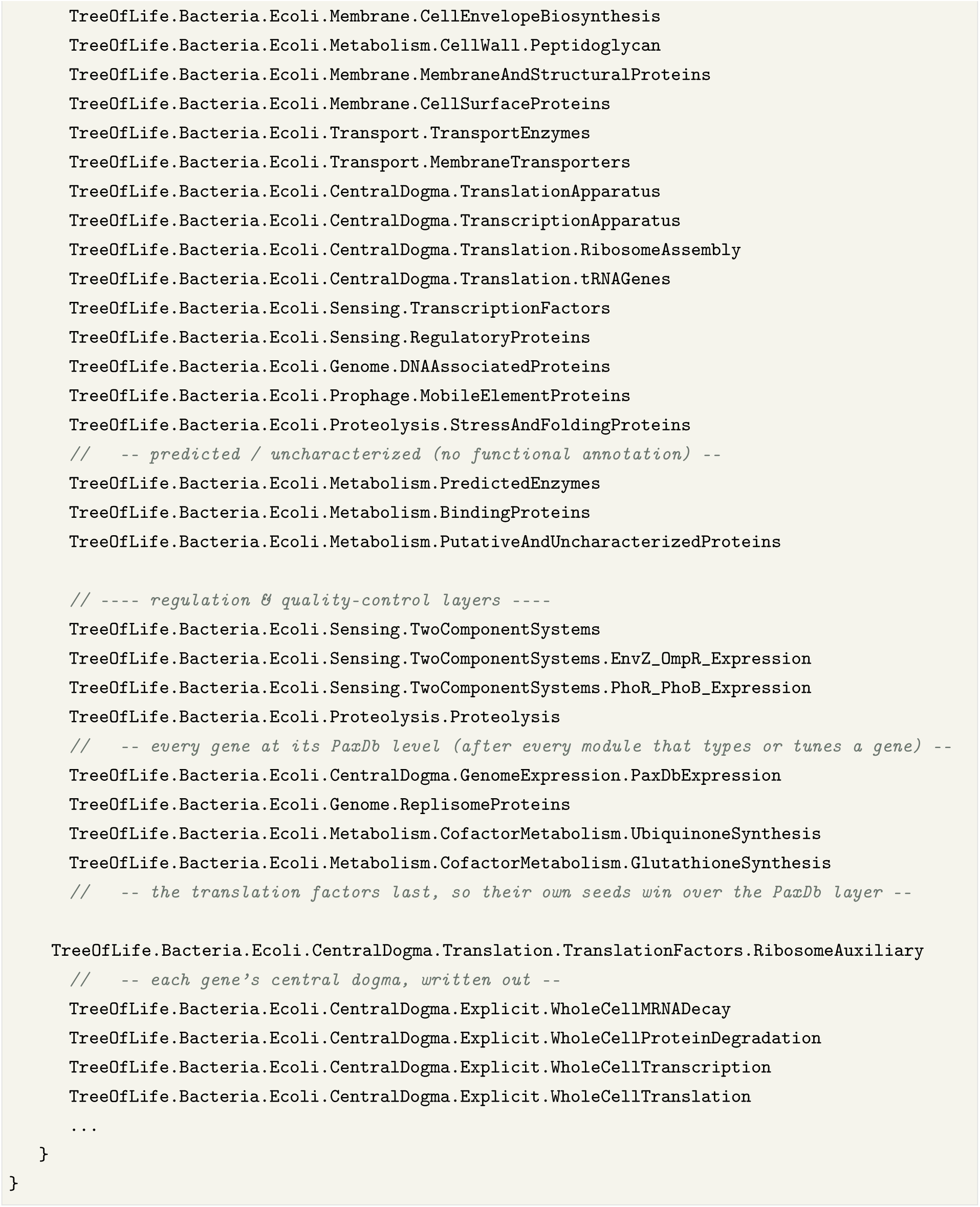

**Listing 1.** The cell of version 1.0 as a composition of modules: a rod born 2 *µ*m long and 0.8 *µ*m wide, whose periplasm and cytoplasm import the model’s modules by their place in the model library: the core of each compartment, the metabolic pathways, the enzymes grouped by EC class, the envelope, transport and central-dogma machinery, the regulators and two-component systems, proteolysis, the expression layer that sets promoter constants toward PaxDb levels, and the four files that write out each gene’s transcription, mRNA decay, translation and protein degradation. Code lines verbatim from the model’s cell definition; comments shortened, and the shared pools of RNA polymerase, sigma factors and ribosomes at the end left out, as are the imports that version 1.0 switches off, among them the SOS response, the pentose phosphate pathway, the glyoxylate shunt and the two prophage modules.

### A genome-wide central-dogma layer

Every gene is expressed, and each gene’s expression is written out on its own, from promoter binding to protein degradation (Listing 2). RegulonDB assigns each gene a primary sigma factor [7], and where it names both *σ* ^70^ and an alternative sigma, the model files the gene under the alternative one by convention, since alternative sigma factors direct adaptive responses [27, 28] (Table 3). In version 1.0 the *σ* ^70^ holoenzyme transcribes nearly every gene, and the alternative sigmas transcribe eight genes written with them: *σ* ^38^ the sensor kinases BaeS, CpxA and EvgS and, with ppGpp bound, the ribosome modulation factor RMF; *σ* ^32^ CreC and PhoQ; *σ* ^54^ ZraS; and *σ* ^28^ CheA. Each of these genes except RMF also keeps a *σ* ^70^ promoter with the same constant, so only RMF depends on an alternative sigma alone, and *σ* ^24^ and *σ* ^19^ transcribe no gene. The core RNA polymerase binds a sigma to form the holoenzyme, the holoenzyme docks the promoter, and in the model the open complex releases the sigma for reuse and loads one lumped complex of the elongation factors NusA, NusG, Rho and GreA. Elongation draws the four NTPs with saturable kinetics, and its capacity follows cell length, a stand-in for the gene dosage that overlapping replication rounds raise through the cycle in fast-growing cells [29]; the model itself replicates one round at a time and keeps a fixed number of copies of each gene per cell. Decay returns each transcript as NMPs. Translation runs on polysomes and consumes charged tRNAs, made by twenty charging reactions at a cost of ATP to AMP and pyrophosphate per charge. Ribosomes assemble from seven-copy rRNA operons and 56 proteins, and subunit assembly requires the GTPases Era, Der and ObgE. The 30S subunit takes the 21 small-subunit proteins, and the 50S subunit takes the 33 large-subunit proteins together with YkgM and RpmJ2 (YkgO), paralogs of L31 and L36. Each protein enters at one copy per subunit, so L7/L12 enters once, where the ribosome of *E. coli* MG1655 carries four copies as two dimers bound to L10 [30]. The cell makes the two paralogs under zinc limitation [31, 32], and each takes the place of its zinc-binding counterpart, L31 or L36, at the same site in a fraction of ribosomes [33, 32]; version 1.0 requires both paralogs in every 50S subunit beside L31 and L36, which makes *ykgM* and *ykgO* necessary for ribosome assembly in the model. These steps abstract the core machinery of bacterial gene expression: promoter binding and initiation [34] and ribosome loading and elongation [35, 36]. The coupling of transcription to translation, in which the leading ribosome sets the elongation rate of RNA polymerase [37, 38], is not represented: elongation depends on the NTPs and on cell length only.

**Table 3.** The seven sigma factors of the model: the role each usually plays in *E. coli* [27, 28, 41], the number of genes for which RegulonDB names it as the primary sigma (Genes; 4,403 genes in all), the genes it transcribes in version 1.0, and how tightly it holds the core RNA polymerase in the model: the dissociation rate constant *k_r_* of its holoenzyme and *K_d_* = *k_r_/k*, with *k* = 10^5^ M*^−^*^1^ s*^−^*^1^ for all seven (model values). Maeda *et al.* measured an apparent *K_d_* of 0.26 nM for the *σ* ^70^ holoenzyme [39]. *σ* ^38^ transcribes RMF only with ppGpp bound, and every gene of the version 1.0 column except RMF also keeps a *σ* ^70^ promoter.

| Sigma | Protein | Usual role | Genes | In version 1.0 | *k<sub>r</sub>* (s<sup>−1</sup>) | *K<sub>d</sub>* (nM) |
|---|---|---|---|---|---|---|
| σ<sup>70</sup> | RpoD (*rpoD*) | housekeeping: most genes of growing cells | 3,540 | nearly every gene | 0.001 | 10 |
| σ<sup>38</sup> | RpoS (*rpoS*) | general stress response and stationary phase | 336 | BaeS, CpxA, EvgS, RMF | 0.02 | 200 |
| σ<sup>32</sup> | RpoH (*rpoH*) | heat shock | 166 | CreC, PhoQ | 0.003 | 30 |
| σ<sup>54</sup> | RpoN (*rpoN*) | nitrogen assimilation and other promoters that need an activator | 144 | ZraS | 0.002 | 20 |
| σ<sup>24</sup> | RpoE (*rpoE*) | envelope stress in the periplasm and outer membrane | 136 | none | 0.01 | 100 |
| σ<sup>28</sup> | FliA (*fliA*) | flagella and chemotaxis | 76 | CheA | 0.04 | 400 |
| σ<sup>19</sup> | FecI (*fecI*) | ferric citrate uptake | 5 | none | 0.06 | 600 |

The seven sigma factors bind one pool of core RNA polymerase by mass action, with the association rate constant *k* = 10^5^ M*^−^*^1^ s*^−^*^1^ for all seven and the dissociation rate constants *k_r_* of Table 3, so the model’s dissociation constants (*K_d_* = *k_r_/k*) run from 10 nM for *σ* ^70^ to 600 nM for *σ* ^19^. The *k_r_* values are model values that encode a rank order of affinity (*σ* ^70^ *> σ* ^54^ *> σ* ^32^ *> σ* ^24^ *> σ* ^38^ *> σ* ^28^ *> σ* ^19^), and the code gives no measured source for them. A comparison of all seven *E. coli* sigma factors on the same core enzyme found *σ* ^70^ bound most tightly, with an apparent *K_d_* of 0.26 nM, followed by *σ* ^54^ and *σ* ^28^, with *σ* ^38^ the weakest [39]. The model shares the two tightest binders with this order and places *σ* ^28^ sixth where the measurement places it third, and its *K_d_*for *σ* ^70^ is about 40 times the measured one. Four promoter constants of the sigma-factor genes were set by hand: the *rpoE*, *rpoN* and *fecI* promoters at a tenth of their earlier value, since these sigmas over-accumulated without active degradation, and the *fliA* promoter at zero, since FliA over-production took the core polymerase from *σ* ^70^. FliA is therefore not made during a run, and its starting amount dilutes with growth. Table 3 lists the seven sigma factors with the roles they usually play, the number of genes for which RegulonDB names each as the primary sigma, the genes each transcribes in version 1.0 and their affinity for the core polymerase in the model; the housekeeping *σ* ^70^ regulon is by far the largest (3,540 genes).

A generated expression layer sets the promoter constants of 2,027 genes toward their PaxDb abundance [40]; in the compiled cell 2,016 of these settings take effect, and 11 genes keep a constant set by hand in a later module. PaxDb abundances are converted at 5.5 nM per ppm. Each constant was set in one step from wild-type runs of an earlier development version: the new constant is the previous one times the PaxDb level divided by the level the protein reached, for the genes more than 1.5-fold from their PaxDb level, with a minimum value for 8 genes read by presence gates (Essential genes and deletions). The other genes keep their class rate. Proteins also start at their PaxDb levels, except where a module sets another seed (GapA in Listing 2), so agreement with PaxDb checks this calibration; during the runs the proteins drift above it (Physiological values).

Each gene’s steps carry the gene’s own composition, read from the GenBank record of K-12 MG1655 (U00096.3) [1, 26]: transcription draws the four NTPs in the counts of the gene’s coding sequence with its stop codon and releases as many pyrophosphates, decay returns the same nucleotides as NMPs, translation spends two GTP per residue and the charged tRNAs of the protein’s own sequence, and degradation returns its amino acids (Listing 2, for *gapA*); selenocysteine counts as cysteine. Of the 4,374 genes with these steps, 4,196 take their own composition. The other 178, insertion-element copies, complexes named as genes, genes named by their locus tag and genes whose record gives no protein sequence, keep one generic composition of 250 of each nucleotide and 300 residues, and the 56 proteins of the ribosome assembly module (the 54 ribosomal proteins and the paralogs YkgM and RpmJ2) keep the rows of that module, which already carry their own sequences. The rate constants are those of the generic composition, so each gene makes as many transcripts and proteins and each copy costs what its sequence costs. Weighted by the flux through each gene in the wild-type run, transcription draws 14% fewer NTPs, since the well-expressed genes are shorter than 1,000 nucleotides, translation draws 1% more residues, and the demand for single amino acids shifts by *−*16% (cysteine, histidine) to +31% (glutamine). Every gene’s transcription and decay, and its synthesis and degradation, return the same nucleotides and amino acids they take. Against the same cell written with the generic composition (seeds 1 and 2 to *t* = 20,000 s), the cell reached its sixteenth division at 19,300 and 18,900 s against 19,000 and 19,200 s, no amount went negative, and the median ratio of each protein’s level was 0.99 and 0.98 over *t* = 1,000 to 3,000 s and 1.00 and 0.99 over 10,000 to 20,000 s, against 1.01 and 1.02 between the two seeds of one build; a run took about 20% longer to compute (7,900 against 6,500 s).

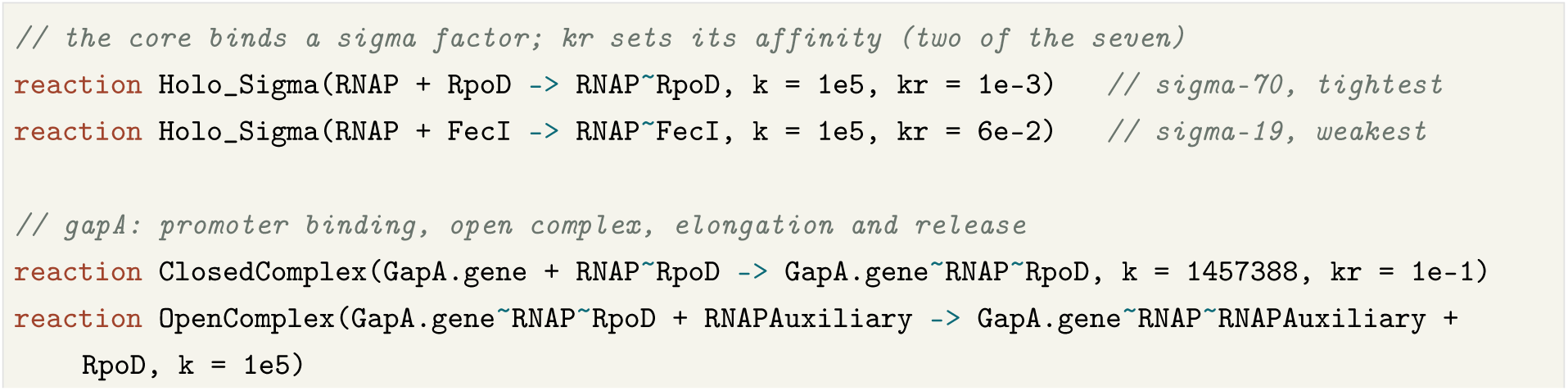
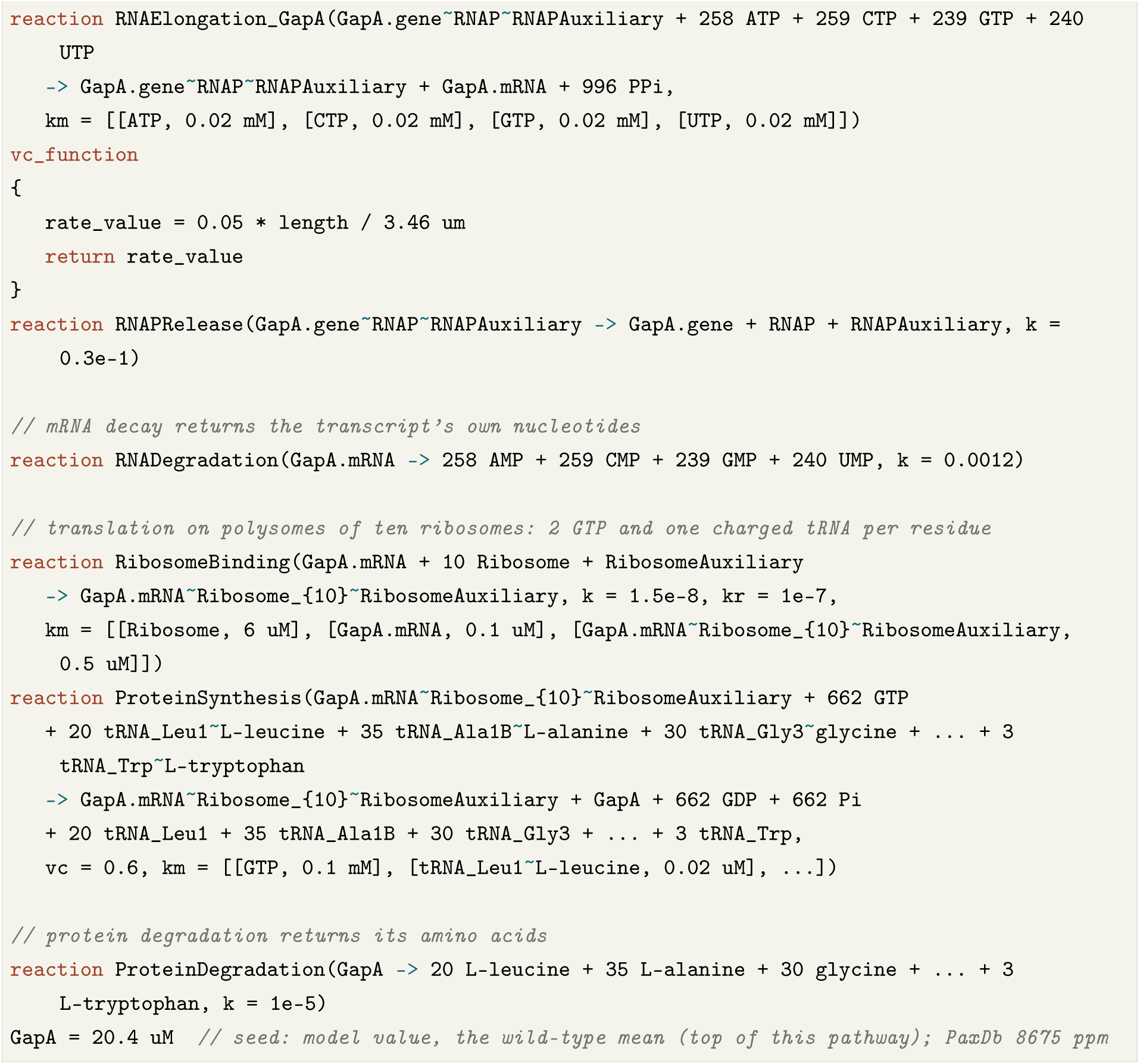

**Listing 2.** One gene written out: *gapA*, glyceraldehyde-3-phosphate dehydrogenase (331 residues), through every step of its central dogma, from promoter binding to protein degradation, after two of the seven sigma holoenzymes. The nucleotides of transcription and decay and the residues and charged tRNAs of translation and degradation are counted from the gene’s coding and protein sequences in the GenBank record U00096.3 (996 nucleotides with the stop codon). Version 1.0 writes every gene’s steps this way, in four files generated from the model’s sigma-class templates (transcription, mRNA decay, translation and protein degradation) that the cell imports after its modules; a step a module writes, such as this gene’s promoter binding and mRNA decay, stays in the module. Code lines verbatim, long lines wrapped and shortened at …; comments ours except the seed line’s. The seed of GapA, 20.4 *µ*M, is the level a wild-type run reaches, a model value; its PaxDb level, 8,675 ppm, would give 47.7 *µ*M.

### An enzyme-kinetics core

Within the genome-wide envelope, roughly 500 metabolites and 1,000 genes carry enzyme kinetics, with constants measured where the model files record a source and model values elsewhere: of the 6,186 reactions outside the per-gene steps of the central dogma, 126 cite a source for some or all of their values in the code, 119 carry marked model values, 5,931 name no source and 10 were not traced (Methods). Central carbon metabolism is written as explicit enzyme reactions with *k*_cat_, *K_m_* and copy number: glycolysis (14 enzymes, 36 reactions), the TCA cycle (17 enzymes, 31 reactions) and oxidative phosphorylation (4 enzymes, 13 reactions), together with ten amino-acid groups and cofactor, nucleotide and lipid metabolism.

Two parts of central carbon metabolism are missing or partial. Version 1.0 does not run the pentose phosphate pathway: its module is written, but the cell does not import it, so no reaction makes 6- phosphogluconolactone, ribulose 5-phosphate, sedoheptulose 7-phosphate or erythrose 4-phosphate. Ribose 5-phosphate, ribulose 5-phosphate and erythrose 4-phosphate are held at fixed values until the pathway runs, and the cell’s NADPH comes from its other NADP-reducing reactions. Glucose enters by three written routes: an ATP-driven transport with no enzyme (*k* = *k_r_* = 0.015, so it also runs backward), phosphotransfer from phosphoenolpyruvate through the mannose permease (ManZ, *k* = 100 s*^−^*^1^, with *K_m_* 50 *µ*M for glucose and for phosphoenolpyruvate), and phosphotransfer through PtsG and MalX at *k* = 0.01, which carries little glucose at their levels. The glucose-specific phosphotransferase system is therefore written only in part, and catabolite control through it is not modeled.

Each enzyme’s level comes from its own expression, with its promoter constant aimed at its measured abundance [40, 15], and turnover numbers measured *in vitro*, which correlate with catalytic rates inferred *in vivo* [42], set its capacity where they are known (Listing 3). Committed steps of amino-acid and nucleotide synthesis run one way under end-product feedback, with measured inhibition constants where they exist (ArgA, SerA; the model files cite their sources), an inhibition constant at the measured pool otherwise (HisG, ProB), or one set so that synthesis meets the demand at the measured pool (PyrB), and uptake from the medium slows as each intracellular pool approaches its measured level [14]. Because each enzyme’s gene is the same object, one protein declaration ties its expression to its catalysis, and deleting or retuning it is a one-line edit.

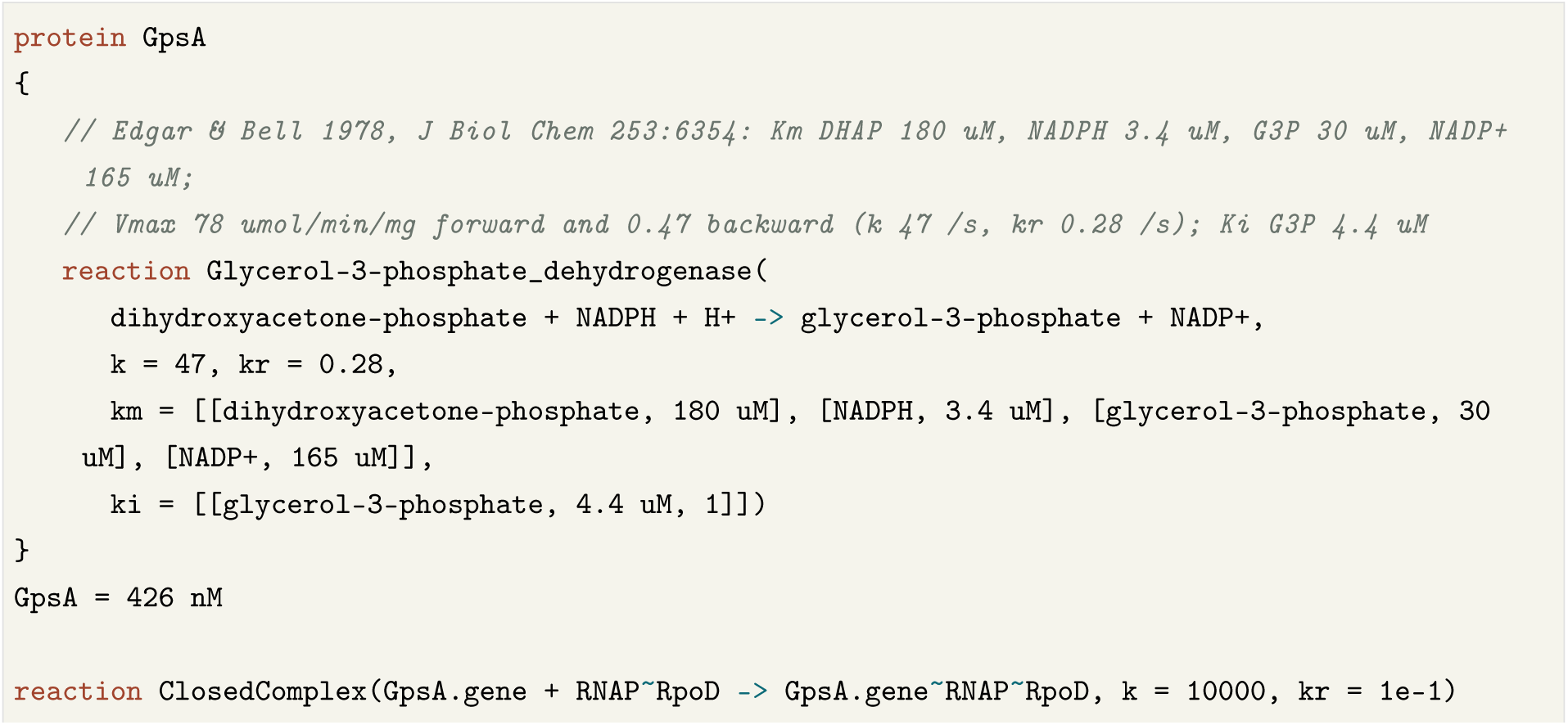

**Listing 3.** An enzyme of the metabolic core with all its constants from one study. Glycerol-3-phosphate dehydrogenase (GpsA) is written in the direction the cell runs it, toward phospholipid, with the constants Edgar and Bell measured for the purified *E. coli* enzyme [43]: *K_m_*for its four substrates and products, turnover in each direction from the measured *V*_max_, and inhibition by its product glycerol 3-phosphate (*K_i_* 4.4 *µ*M), which acts at a regulatory site. Its abundance and its promoter constant are model values aimed at its PaxDb level. From the model’s glycolysis module; source comments shortened.

### How close the virtual cell is to *E. coli*

A life scientist will first ask how closely the virtual cell behaves like *E. coli*. Table 4 brings together the main comparisons with measurement in this paper, each with the measurement it rests on and the version it ran on; its fourth column gives the model-to-measured ratio or the share that agrees, and its last column says how the model came to agree. Supplementary Tables S1 to S3 give the essential-gene candidates, the measured pools and the held pools one by one. Much of the agreement is built in. The replication fork rate was chosen so that a round fits a 20-minute cycle; ATP is capped at its measured pool, and 43 of the 95 compared metabolite pools start within 10% of their measured values; the promoter constants were set toward PaxDb abundances; the presence gates, which read 154 essential gene products, were written from the calls of the essential-gene screens; and the constants of the two tested two-component systems were set from the responses they are compared with. Rows of these kinds show that the whole cell keeps what was put into it, and the rows marked *none* test the model.

**Table 4.** How close the virtual *E. coli* is to measured *E. coli*. Values of *E. coli* version 1.0 (seeds 1 and 2; pools over *t* = 1,000 to 3,000 s, proteins over that window and *t* = 10,000 to 20,000 s, ribosomes after *t* = 8,000 s) unless a development version (B to E, Methods) is named. Basis says how the model came to agree: *calibrated*, a model value was chosen to reproduce this measurement or the data behind it; *seeded*, the run starts at the measured value; *capped*, an overflow reaction bounds the value (ATP); *cap not reached*, a rule degrades free ribosomes above 16 *µ*M, and the free pool of version 1.0 stays below it (mean 12 *µ*M); *wired*, a rule written from the same data gives the result; *coded size*, the value follows from the cell dimensions written in the model; *mixed*, rows of several kinds; *none*, the model was not set to agree, so the row tests it. Of the 95 pools, coenzyme A and *S*-adenosylmethionine are held at their measured values, 6-phosphogluconate is not diluted by growth, two of the three pentose phosphates summed as one pool are held, 43 start within 10% of their measured value, and the uptake of 16 amino-acid pools from the medium slows at their measured values.

| Property | Virtual *E. coli* | Measured *E. coli* | Agreement | Basis |
|---|---|---|---|---|
| **Growth and size** |  |  |  |  |
| Doubling time in rich medium | 18–23 min | 22 min in LB [15] | 0.8–1.1 | calibrated |
| Ribosomes per cell | 62,000–63,000 | about 73,000 at a 20-min doubling [29] | 0.86 | cap not reached |
| Cell volume, version C | 1.63 fL | 4.29 fL in LB, estimated from growth rate [15, 44] | 0.38 | coded size |
| **Metabolome** |  | 95 of 103 absolute pools [14] |  |  |
| Pools within 2-, 4- and 10-fold at *t* = 0 | 56, 61 and 78 |  | 59%, 64%, 82% | seeded |
| Pools within 2-, 4- and 10-fold, *t* = 1,000 to 3,000 s | 41, 67 and 82 |  | 43%, 71%, 86% | mixed |
| Pools within 2-, 4- and 10-fold, *t* = 10,000 to 20,000 s | 37, 52 and 65 |  | 39%, 55%, 68% | mixed |
| ATP | 9.60 mM | 9.6 mM | 1.0 | capped |
| GTP | 6.17 mM | 4.9 mM | 1.3 | seeded |
| NAD<sup>+</sup> | 3.03 mM | 2.6 mM | 1.2 | seeded |
| Phosphoenolpyruvate | 0.23 mM | 0.18 mM | 1.3 | none |
| NADH | 0.44 mM | 0.083 mM | 5.3 | none |
| NADP<sup>+</sup> | 1.30 mM | 2.1 *µ*M | 620 | none |
| L-aspartate | 8.0 mM | 4.2 mM | 1.9 | none |
| L-glutamate | 10.6 mM | 96 mM | 0.11 | none |
| **Proteome** (with complexes) |  |  |  |  |
| Median against the LB proteome | 5.9–6.2 times | [15] | 58–60% in 10-fold | none |
| Median against PaxDb | 2.4–3.5 times | [40] | 68–75% in 10-fold | seeded, calibrated |
| **Essential genes** |  | Keio, PEC and TraDIS [45] |  |  |
| Wired to a growth process, of the genes two or three screens call essential (count from the code) | 293 of 293 |  | 100% | wired |
| Deletion stops or slows growth, version B | 104 of 139 tested |  | 75% | mixed |
| gene read by a presence gate | 91 of 91 |  | 100% | wired |
| gene acting through its own product | 13 of 48 |  | 27% | none |
| **Responses to the medium**, versions D and E |  |  |  |  |
| OmpR∼P after NaCl 50 to 350 mM | 5.5% to 34% | 5% and 27% (20% sucrose, taken as 0.3 M NaCl) [46] |  | calibrated |
| *ompC* rise (model: mRNA; measured: *ompC-lac* expression rate) | 2.0-fold | 3.4-fold until about 40 min, then 2-fold [47] | 0.6 (1.0 against the later rate) | calibrated |
| *ompF* fall (model: mRNA; measured: *ompF-lac* expression) | 0.46 | 0.5 [47] | 0.9 | calibrated |
| PhoB after phosphate falls to 1 *µ*M (model: PhoB∼P; measured: total PhoB) | 2.8 *µ*M at 20–50 min, *phnC* last | half-rise about 25 min [48], *phnC* last [49] | same order | calibrated |

Growth is close: the cell doubles every 18 to 23 minutes, against 22 minutes for LB-grown cells [15], a calibrated agreement. Of the 103 metabolites whose absolute concentrations were measured [14], the model holds 95; 41 of these lie within twofold and 82 within tenfold over *t* = 1,000 to 3,000 s, against 56 and 78 at *t* = 0. NAD^+^ and phosphoenolpyruvate run at 1.2 and 1.3 times their measured levels, NAD^+^ from a start at its measured level and phosphoenolpyruvate from a start at about twice its pool, while NADH, NADP^+^, glutamate and aspartate show where the metabolism still departs. Proteins run about six times above the LB proteome at the median, in a cell whose ribosome count is about 86% of the measured count and whose volume on development version C was 38% of the volume estimated for LB-grown cells. On version B, 104 of the 139 tested genes that two or three screens call essential stop or slow growth when deleted: all 91 that a presence gate reads, and 13 of the 48 that act through their own product. On versions D and E the osmolarity and phosphate sensors respond in the measured direction and promoter order, which their constants were set to give.

### Physiological values

A virtual cell is judged first by its physiological values, the quantities measured in growing cells: how fast it divides, how large it is, how many ribosomes and other proteins it holds and at what concentrations it keeps its metabolites. The metabolite reference is the absolute metabolome of *E. coli* K-12 grown on glucose minimal medium [14], 103 metabolites spanning nearly six orders of magnitude, 95 of which the model holds; the protein references are the LB proteome of Schmidt *et al.* [15] and PaxDb [40], and the ribosome reference is the counts of Bremer and Dennis [29]. The simulated cell grows in a rich medium, so pools that the medium supplies are expected to differ. Each pool is averaged over a window of seeds 1 and 2 and scored by its log_2_ fold change from the measurement, and the model is scored by the number of pools within 2-, 4- and 10-fold (Methods).

A run starts from one cell in a 10 *×* 10 *×* 1 cm dish of a modified MOPS EZ Rich defined medium [50]. The model medium has the recipe’s 19 amino acids and four bases at the recipe’s concentrations and its main salts, with four departures: glucose is 4.1 mM (the recipe’s 0.2% is 11.1 mM), glutamate is 11 mM (the recipe has 0.6 mM), the vitamins are a different set, and iron is 10 nM (the recipe has 10 *µ*M). H^+^,

O_2_ and CO_2_ are held at 1.2 mM (Methods). The simulation runs at resolution level 6, in ticks of 2*^−^*^6^ s (Methods). Cell division fires when replication has finished a second genome copy, the cell has reached 4 *µ*m, its wall is intact and the divisome is present, and elongation is driven by the cell’s protein.

The replication fork rate is a model value chosen so that a round fits a 20-minute cycle: two forks of 2,272 nucleotides per second at full rate copy the model’s genome of 5.0 Mb (the sequenced genome has 4.64 Mb) in 1,100 s. Forks in *E. coli* B/r at 37 *^◦^*C run less than half as fast: their C period, the time to replicate the chromosome, is about 40 minutes at doubling times of 20 to 60 minutes, and the D period from the end of replication to division about 20 minutes [51]. Each round of the model starts as the cell divides, from an origin held loaded, and the model runs one round at a time. It has neither the overlapping rounds of fast-growing cells nor initiation controlled by DnaA, and it has no D period, since division can fire as soon as a round ends. The elongation set point was tuned so that the cell reaches 4 *µ*m together with replication at about 1,100 s. The cycle is therefore set mainly by the replication rate, and the doubling time is a calibrated value.

Values are brought toward measurement pathway by pathway. In such a pathway each enzyme takes its measured turnover number, Michaelis constants and level where they exist, and a model value, marked in the code, where none is measured or where the measured enzyme cannot carry the flux of a cell that divides every 20 minutes: at its measured level and turnover, nicotinate-mononucleotide adenylyltransferase (NadD) makes a twelfth of the NAD that growth dilutes, so its level is a model value. Protein levels start at their PaxDb levels unless a module sets another seed, and 2,016 promoter constants are set toward PaxDb levels. A pathway brought to measured values joins the model as one unit, after the same whole-cell test as a released pool (Methods). The units applied so far put the cell-wall pathway on measured kinetics and the replisome at measured levels; make FAD, ubiquinone, nine cofactors, glutathione, gluconate and cyclic AMP by their pathways, and NAD through NadD and NadE; and run asparagine synthetase (AsnA), nicotinate phosphoribosyltransferase (PncB) and phosphoenolpyruvate carboxylase (Ppc) one way, Ppc at its measured turnover number.

Some agreement is built in and is not a prediction. Besides the doubling time, an overflow reaction burns ATP above 9.6 mM, the measured pool, so the ATP record tests only whether the supply of ATP keeps up with demand. Coenzyme A, *S*-adenosylmethionine and two of the three pentose phosphates that the comparison sums as one pool are held at fixed values, and 6-phosphogluconate keeps its concentration as the cell grows (next section). The feedback and uptake constants of lysine, serine and the aromatic amino acids are set at their measured pools; in all, uptake from the medium slows at the measured pool for 16 of the compared amino-acid pools. Of the 95 pools, 43 start within 10% of their measured value, and at *t* = 0, before any reaction has run, 56, 61 and 78 lie within 2-, 4- and 10-fold.

On *E. coli* version 1.0, the model this article describes (seeds 1 and 2 to *t*= 20,000 s), the cell divides every 1,100 to 1,400 s through 16 divisions with no negative-concentration event, near the 22-minute doubling of the LB-grown cells whose proteome is the reference [15] and the 20-minute doubling of *E. coli* B/r in LB with glucose at 37 *^◦^*C, the fastest in the tables of Bremer and Dennis [29]. The absence of negativeconcentration events reflects the redraws and the depletion guard of the simulator as well as the model (Methods). ATP averages 9.58 and 9.49 mM over *t* = 1,000 to 20,000 s. In seed 1 it stays at or above 9.51 mM in every 100 s snapshot; in seed 2 it falls below 9 mM once, from *t* = 14,300 to 14,800 s, with a lowest point of 5.75 mM at *t* = 14,600 s. Counting free ribosomes and those in polysomes and other complexes, the cell holds on average 62,400 and 62,700 ribosomes after *t* = 8,000 s, about 86% of the 73,000 per cell measured at a 20-minute doubling [29]; its free ribosomes average 12 *µ*M, below the 16 *µ*M above which a rule degrades them. Of the 95 measured pools, 56, 61 and 78 lie within 2-, 4- and 10-fold at *t* = 0, 41, 67 and 82 over *t* = 1,000 to 3,000 s (Figure 2) and 37, 52 and 65 over *t* = 10,000 to 20,000 s. Of the 17 pools that leave tenfold between the two windows, ten fall as fast as growth dilutes them: no reaction makes glycerate, 2,3-dihydroxybenzoate or 6-phosphogluconolactone, and the other seven gain almost nothing from the reactions written to make them. Three fall more slowly, and four (shikimate, quinolinate, dGMP and glycerol 3-phosphate) change by less than twofold and cross the line. NAD^+^ and phosphoenolpyruvate run at 1.2 and 1.3 times their measured pools (Table 4). NADH falls from its 1 mM start to about five times its measured pool. NADP^+^ stays near its 1.2 mM start, about 620 times its measured pool, because NAD kinase holds it at about half of NAD^+^ (below). L-glutamate stays near the 11 mM of the medium, a ninth of its measured pool, and aspartate runs at about twice its pool, since the measured inhibition of Ppc by aspartate is not yet written. The pool-by-pool comparison for all 95 metabolites is Supplementary Table S2.

**Figure 2.**
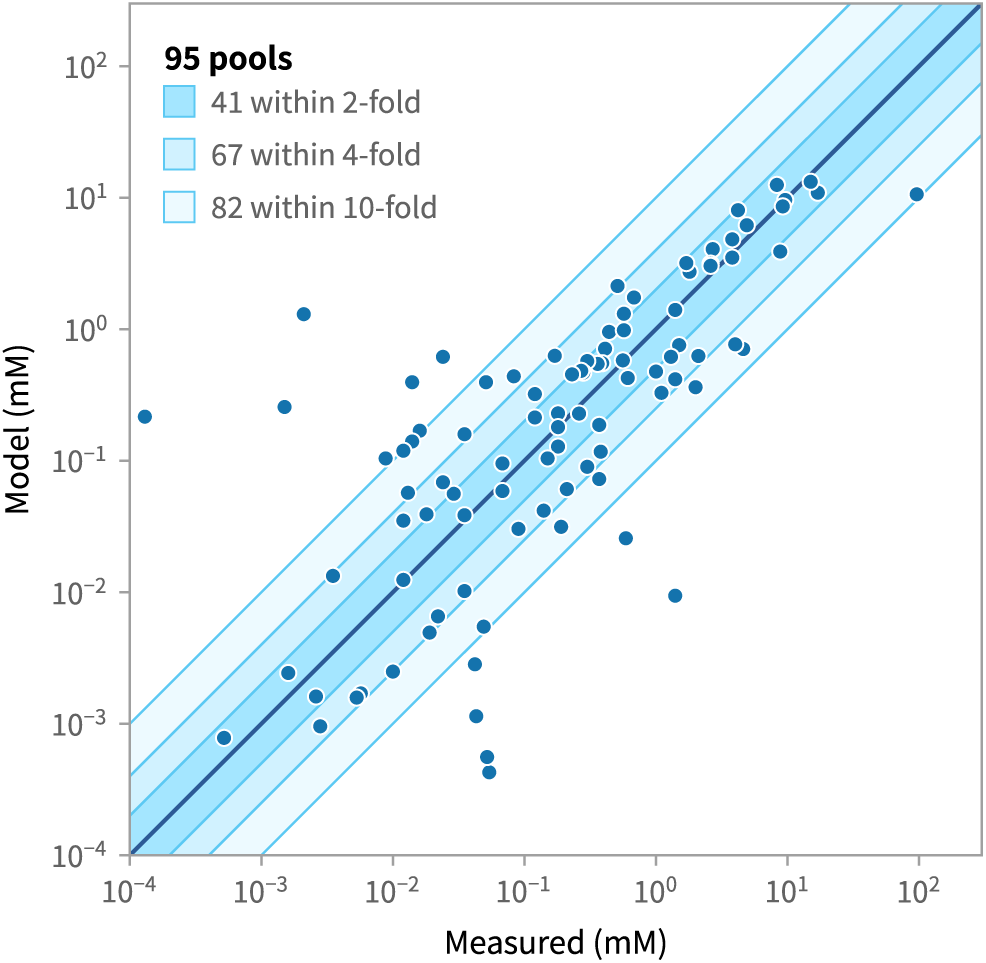
Model against measurement for the 95 metabolite pools with absolute concentrations in glucose-grown *E. coli* [14]: means over *t* = 1,000 to 3,000 s of seeds 1 and 2 on *E. coli* version 1.0. Lines and shading mark equality and 2-, 4- and 10-fold, and the key counts the pools within each band. The pools include coenzyme A and *S*-adenosylmethionine, held at their measured values, ATP, capped at its measured value, and 43 pools that start within 10% of their measured values (Table 4).

Thirteen of the 95 pools lie outside tenfold over *t* = 1,000 to 3,000 s, and each departure traces to a choice written in version 1.0. The causes below are read from the code, and the pools in the runs sit at or near the ratios the code sets. Six of the thirteen start at values set 10 to 1,000 times above their measured values, which the model file keeps as comments beside them: adenosine, NADP^+^, adenine, GMP, dAMP and dATP. The reactions that tie each of them to a neighboring pool are written reversible with equal forward and backward constants and with Michaelis constants within 3% of the starting values, so each comes to rest with the two pools in the ratio of their starting values (Methods). Adenosine kinase holds adenosine at 0.46 times AMP (measured ratio 0.0005), NAD kinase holds NADP^+^ near 0.49 times NAD^+^ (0.43 in the runs; measured 0.0008), adenine phosphoribosyltransferase holds adenine near 0.53 times AMP, guanylate kinase holds GMP at 0.35 times GDP (measured 0.035), and two kinase steps hold dAMP and dATP at 0.31 and 0.50 times dADP, which starts at 100 times its measured pool. Quinolinate and shikimate sit inside chains of such reactions and stay near starting values set at ten times their measured pools, just inside tenfold. The rich medium supplies two pools that the glucose minimal medium of Bennett *et al.* lacks: cytosine, which the permease CodB takes up one way and only a reaction at the shared placeholder constant uses, and adenine, which also enters through reversible permeases. Gluconate starts at its measured 42 *µ*M, but its three permeases (GntP, GntT and GntU) run both ways at equal forward and backward constants against a periplasm that holds no gluconate, so it leaks out and settles near 3 *µ*M, where the dephosphorylation of 6-phosphogluconate by YieH and YqaB balances gluconate kinase (GntK) and the efflux. Dihydroorotase holds carbamoyl-aspartate at about twice dihydroorotate (measured ratio 49), and ornithine carbamoyltransferase holds citrulline at no more than 5.2 times ornithine (measured 140), since carbamoyl phosphate, which drives this step in the cell, has no Michaelis constant in the reaction and does not enter its rate. *N*-acetylornithine runs low because arginine from the medium inhibits the first enzyme of its pathway, *N*-acetylglutamate synthase, at the measured inhibition constant, and the rate constant of that enzyme is a model value set to leave about half of the arginine demand to uptake. FMN and 4-hydroxybenzoate are used by enzymes that run far below their Michaelis constants: the riboflavin kinase and FAD synthetase RibF, at 21 and 120 times its measured turnover numbers (model values), keeps FMN near 0.4 *µ*M while FAD runs at about four times its measured pool; the measured product inhibition of chorismate lyase (UbiC) and a Michaelis constant of 2 *µ*M for UbiA (a model value) hold 4-hydroxybenzoate near 0.6 *µ*M. Which of two likely causes dominates is not tested for adenine (uptake or its ratio to AMP) and for *N*-acetylornithine (inhibition by arginine or the low glutamate pool), and the pyrimidines of the medium may lower carbamoyl-aspartate further by holding CTP and UTP above the levels at which aspartate carbamoyltransferase meets the demand (likely, not tested).

Proteins counted with their complexes, averaged over seeds 1 and 2, run at a median 5.88 times the LB proteome [15] over *t* = 1,000 to 3,000 s and 6.22 times over *t* = 10,000 to 20,000 s, with 60% and 58% of them within tenfold, and at a median 3.51 and 2.36 times PaxDb [40], with 68% and 75% within tenfold. The proteins of the measured set total 7.83 and 7.34 mM in the two windows, against 3.09 mM measured for the same proteins. Since the proteins start at their PaxDb levels, apart from the seeds that modules set, and the promoter constants aim at them, the PaxDb comparison shows how far the proteins drift from their calibration. Part of the excess is set by the targets: the PaxDb levels that the expression layer aims at sit at a median 3.3 times the LB proteome. Ribosomal proteins run at a median 4.9 and 4.6 times the LB proteome in the two windows, and the most abundant other proteins run low: EF-Tu falls from its starting level to 12 *µ*M over *t* = 10,000 to 20,000 s, against 172 *µ*M measured [15], since one gene makes at most about 0.05 transcripts per second, about 2 *µ*M of protein in a 1,200 s cycle. On development version C, the cell’s volume of 1.63 fL was below the 4.29 fL estimated for LB-grown cells [15, 44], so per-cell totals sat closer to measurement than concentrations did. The volume follows from the size written in the model, a rod 0.8 *µ*m wide born 2 *µ*m long that divides at 4 *µ*m.

Figure 3 follows two seeds of development version A, built before the tRNA supply fix and the peptidoglycan wall, to *t* = 40,000 s. The cells pass through 35 consecutive divisions to 2^35^ with one negativeconcentration event (*−*1.2 *×* 10*^−^*^10^ M at *t* = 13,407 s in seed 1), ATP averages 9.56 and 9.57 mM, GTP has a median of 4.6 and 4.8 mM against the measured 4.9 mM [14], and ribosomes average 56,000 and 59,000 per cell after *t* = 8,000 s, against about 73,000 per cell at a 20-minute doubling in the tables of Bremer and Dennis [29]; ATP is capped at its measured pool, and a rule degrades free ribosomes above 16 *µ*M. ATP fell after some divisions on that version: across six runs (seeds 1 to 6), the adenine nucleotide pool dropped by 0.8 mM or more between 100 s snapshots in 9 episodes, each starting within 1,000 s of a division, and ATP was below 6 mM in 6 of 2,346 snapshots. Seeds 1 and 2, shown in the figure, are two of the three runs whose ATP stayed above 6.8 mM after *t* = 1,000 s; in seeds 3, 4 and 6 it fell to 4.4, 5.2 and 5.95 mM.

**Figure 3.**
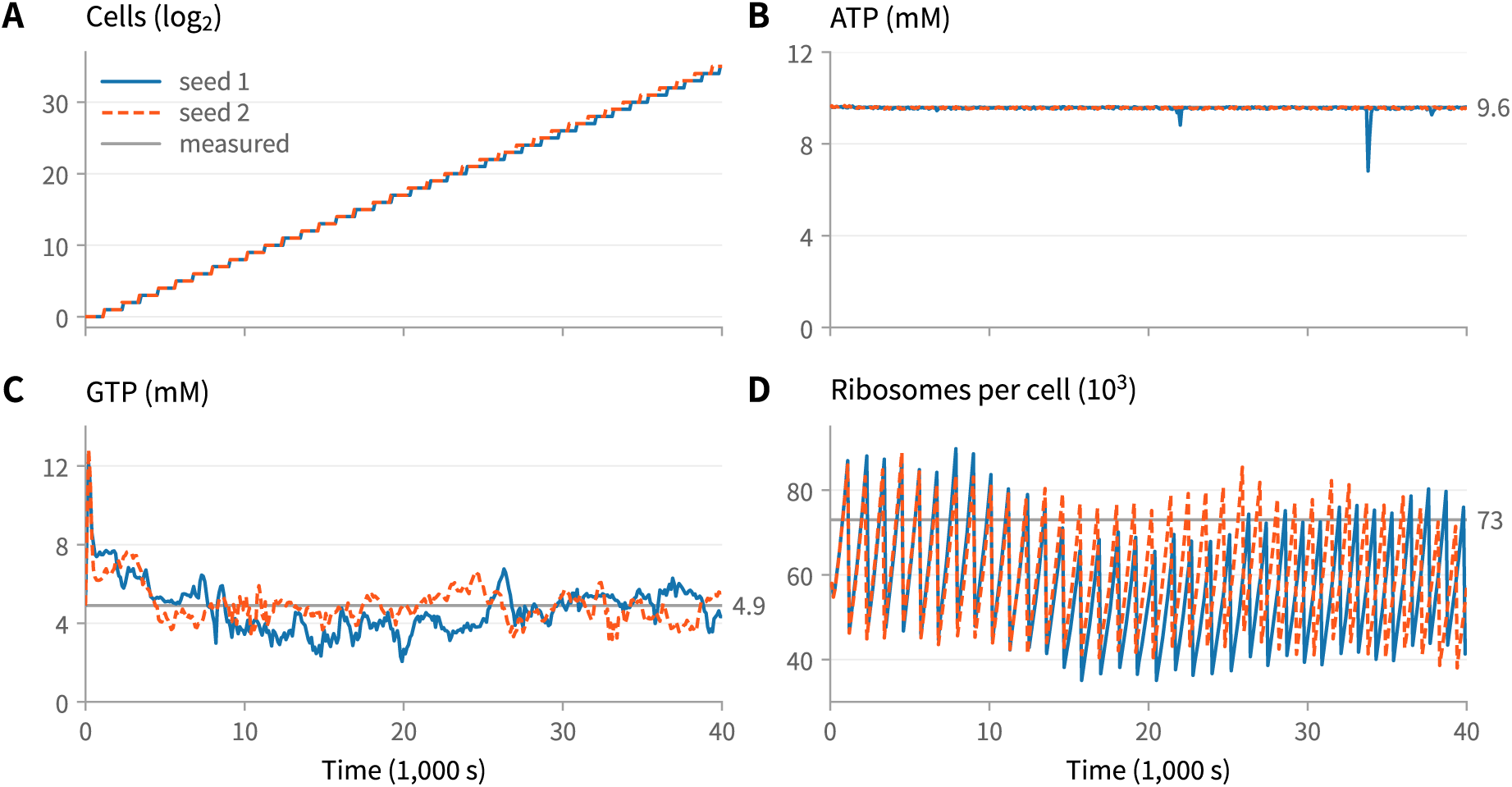
Whole-cell runs to *t* = 40,000 s on development version A, built before the tRNA supply fix and the peptidoglycan cell wall: seeds 1 and 2 of the six that ran. (A) Cells on a log_2_ scale, the count of identical cells that one simulated cell stands for. (B) Cytoplasmic ATP. (C) Cytoplasmic GTP. (D) Ribosomes per cell. Gray lines mark the measured values: ATP 9.6 mM and GTP 4.9 mM in glucose minimal medium [14], and about 73,000 ribosomes per cell at a 20-minute doubling [29]. Two of the plotted quantities are capped: an overflow reaction burns ATP above 9.6 mM, so the gray ATP line is also the cap, and a rule degrades free ribosomes above 16 *µ*M. Snapshots every 100 s, simulated with lpps at resolution level 6.

### Clamped pools and their release

A model of this size is built in stages. A pool whose synthesis is not yet written is clamped: L++ holds it at a fixed value for the whole run (X[:] = value), so the reactions that make or use it have no net effect on it, and the rest of the cell can be tested around it. Each clamp is then released: the pathway that makes the pool is written, on measured kinetics where they exist, each reaction that consumes the pool is given a Michaelis constant for it, and the change joins the model only after the whole cell runs with it on two seeds, together with a set of regression deletions, with no amount driven below zero (Methods). Released so far are pyrophosphate, formate, 5*^′^*-deoxyadenosine, 4-amino-4-deoxychorismate, glycolaldehyde and 4-methylphenol; sulfite, now made by the sulfur assimilation pathway on measured kinetics; the thioredoxin pair, whose split TrxB now sets while its total keeps its concentration as the cell grows; 1-deoxy-D- xylulose 5-phosphate, made by Dxs on measured kinetics; the glycerol-3-phosphate dehydrogenase GpsA, which had been held at zero and now runs on measured kinetics [43]; and, after version 1.0, carbamoyl phosphate under pyrimidine control and the ThiS sulfur-carrier pair.

In version 1.0, 31 pools are clamped, 25 of them in the cytoplasm (Supplementary Table S3): H^+^, O_2_, CO_2_, phosphate (23.9 mM), bicarbonate, ammonium, coenzyme A, ribose 5-phosphate, ribulose 5-phosphate, erythrose 4-phosphate, carbamoyl phosphate, pyridoxal and pyridoxamine phosphate, *S*- adenosylmethionine, the ThiS sulfur-carrier pair, the five folates (tetrahydrofolate, 5,10-methylene-, 10- formyland 5-methyltetrahydrofolate, and dihydrofolate), the replication origin’s loaded polymerase, and three pools that feed the cell wall, *meso*-diaminopimelate and the isoprenoid precursors farnesyl and isopentenyl diphosphate. The other six are H^+^, O_2_ and CO_2_ in the periplasm and in the medium. Ten are clamped by design: H^+^, O_2_ and CO_2_ in the cytoplasm, the loaded origin and the six boundary pools; phosphate is held at its measured level. H^+^ and phosphate take part in hundreds of reactions, and without a held H^+^ pool the reactions that consume protons exhaust it within a second of simulated time. H^+^, O_2_ and CO_2_ are held at 1.2 mM in all three compartments as supply pools for the reactions that use them. Read as a free proton concentration, 1.2 mM would be pH 2.9, so the H^+^ pool stands for no pH, and the model has no pH variable; O_2_ at 1.2 mM is about six times its level in air-saturated water at 37 *^◦^*C, and the gases cross the membrane faster than one tick of the simulation resolves. The loaded origin is a structure the replication fork reads, and the boundary pools stand for the medium. The other 20 stand in for synthesis not yet written. Three more pools are written so that growth does not dilute them: 6-phosphogluconate, which the pentose phosphate pathway would use, and the two forms of thioredoxin.

A clamped pool is right by construction, and a clamp can hide what its missing synthesis would show. With carbamoyl phosphate clamped, carbamoyl phosphate synthetase ran backward and made 3 to 4% of the cell’s ATP from the held pool; with coenzyme A clamped, bursts of pantoate synthesis drew 18 to 27 *µ*M/s of adenine nucleotide into it, where growth needs 0.84 *µ*M/s; and bicarbonate has no maker, because carbonic anhydrase writes the same chemical under another name. Releasing a pool exposes such gaps. Freed folates showed that the cell’s dihydrofolate reductase activity sat on a minor enzyme and that every step of *de novo* purine synthesis ran within a few percent of equilibrium: the cell grew 6 to 7% slower and ATP fell to 2 to 3 mM, and the release was taken back. Purine end-product control, written next, passed its test in one of the two fast-species settings the simulations use and failed in the standard one, and was taken back too; since then a release that touches purine or folate metabolism is tested in both settings. *S*-adenosylmethionine rose from 0.18 to 16 mM when its synthase ran at measured kinetics, because the model lacks the cell’s methylation and polyamine sinks, and it stays clamped until they are written. The pentose phosphate pathway, which releases the three pentose phosphates, and the first step of coenzyme A synthesis are in test; bicarbonate, the vitamin B_6_ phosphates, ammonium, the isoprenoid precursors and *meso*-diaminopimelate follow, and the five folates come off once purine synthesis runs under end-product control in both settings.

### Essential genes and deletions

Each essential process asks for the gene products that carry it out. Replication runs only while the replisome is present (GyrA, GyrB, DnaA, DnaB, DnaC, DnaE, DnaG, DnaN, DnaX and Ssb), division needs the divisome and MukBEF, cell-wall growth needs the peptidoglycan pathway and the other essential envelope machinery, subunit assembly needs Era, Der and ObgE, and translation uses its factors under their gene-product names. The Keio collection deleted every non-essential gene one at a time and recorded those that could not be deleted [52]; PEC (Profiling of the *E. coli* Chromosome) and the transposoninsertion screen TraDIS reach the essential set by other means, and Goodall *et al.* give each gene’s call under all three [45]. The model keeps a list of 377 candidate genes for essential processes, each with its call in that comparison: 248 are essential in all three screens, 45 in two, 46 in one and 38 in none.

A candidate counts as wired when a process the cell needs for growth reads its product, by one of two kinds of route (Table 5). In the first, the model reads the presence of the product itself: a presence gate stops a process while the product is below 10 nM (0.3 *µ*M for FtsZ), and a coded slowdown cuts a rate to a set fraction, half for the replication fork and a fifth for ribosome assembly, a tRNA charging reaction or pyruvate and 2-oxoglutarate dehydrogenase. These gates and slowdowns were written from the screens’ calls, so the requirement they create follows the screens by construction. Six of the gates (SpoT, CohE, WzyE, CydA, CydC and YceQ) stand for deletion phenotypes whose causes the model lacks and represent no function of the protein. In the second kind of route, the product does its own job: it is a subunit consumed as a complex assembles, the only charging enzyme of a tRNA, or the enzyme of a reaction whose product runs out without it. In version 1.0, presence gates read 154 gene products, all of them among the 293 that at least two screens call essential: 16 replisome parts in the rate of the replication fork, 17 divisome and segregation proteins in division, 102 envelope, cofactor and stand-in proteins in cellwall synthesis, 7 tRNA maturation enzymes in tRNA synthesis, 8 ribosome assembly factors in subunit assembly and 4 subunits or modification enzymes in tRNA charging. Coded slowdowns read 23 more. Counted from the code, 360 of the 377 candidates are wired, including all 293 that at least two screens call essential, 155 of them through a gate or a slowdown and 138 through their own function; of all 377, 177 are wired by construction and 183 through their own function. The 67 wired candidates that at most one screen calls essential are expected false positives: the model would score their deletions as harmful where at most one screen does. The 17 with no route are all called essential by at most one screen; for some of them the code gives that as the reason, and the others make products the model holds as fixed pools or have deletion phenotypes that act through mechanisms the model lacks. Among them are *polA*, *priA*, *recA* and *rnhA*, whose coded slowdowns version 1.0 removed, since their products act on a fork only when it stalls or collapses, which the model does not represent. The model writes lysyl-tRNA charging on LysS alone: LysU, the second lysyl-tRNA synthetase of *E. coli*, is expressed but carries only diadenosine tetraphosphate synthesis, so the model counts LysS as the only charging enzyme of its tRNA, a gap in the model.

**Table 5.**
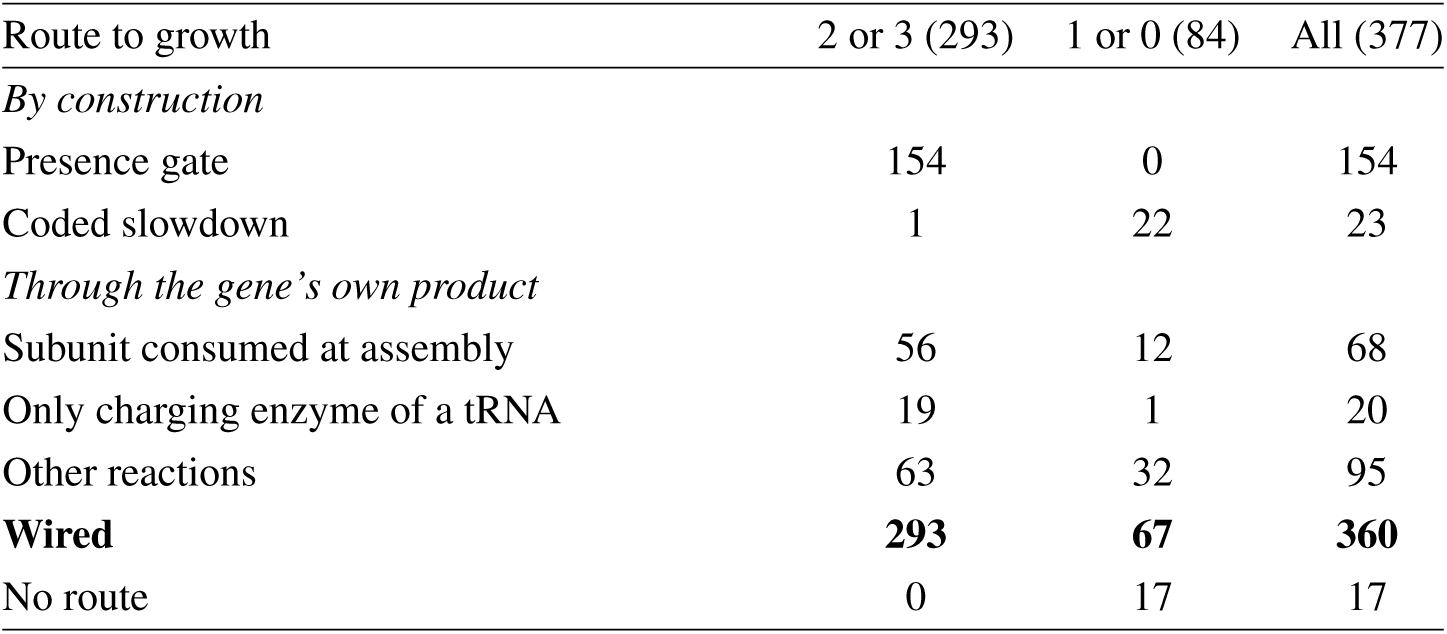
How the model’s 377 essential-gene candidates reach growth in version 1.0, counted from the code, by the number of the Keio, PEC and TraDIS screens that call each gene essential in Goodall *et al.* [45]. Presence gates and coded slowdowns were written from the screens’ calls, so agreement through them holds by construction; the other routes run through the gene’s own product.

A deletion removes a gene’s transcription and its protein at *t* = 0. In a screen of 189 deletions on development version B (seed 1; Methods), 139 genes that at least two screens call essential ran. Scored by the cell count against the wild type at the same time, 86 stopped at one cell, 18 slowed and 35 grew, so 104 of the 139 (75%) stopped or slowed. All 91 tested genes read by a presence gate stopped or slowed, as the gate requires. Of the 48 tested genes that act through their own function, 13 stopped or slowed and 35 grew, and 19 of those that grew make products that version held as fixed pools (coenzyme A, folates, NAD, flavins, thiamine, biotin). Four deletions (*rpsA*, *ribE*, *gapA* and *fbaA*) were recorded as slowed while ending with the wild-type count; counted as slowed, they give 108 of 139. The *adk* deletion, counted as a stop, ran with 121,389 negative-concentration events. Of the 50 tested genes that at most one screen calls essential, 17 stopped or slowed under the same rule, so the screen also scored about a third of these as harmful; four of them (*polA*, *priA*, *recA* and *rnhA*) slowed through the coded slowdowns that version 1.0 removed. In representative deletions on development version A (seed 1 to *t* = 3,000 s), cells lacking FtsZ, GyrA, LpxC, ValS or ThyA stop at one cell, cells lacking RpsC or Era divide once while the ribosomes they already have dilute, and deletions of *argA* and *purF*, whose products the rich medium supplies, grow with the wild type. The *pyrB* deletion first divides at 2,700 s against 1,200 s for the wild type. None of the three screens calls *pyrB* essential, and the 0.2 mM uracil of the medium, which the model takes up and converts to UMP through uracil phosphoribosyltransferase (Upp), should rescue the deletion; the delay is a discrepancy, and on other development versions the same deletion grew with the wild type or first divided at 1,800 to 2,900 s. Each candidate’s screen calls, its route as counted on development version B and, for the 189 deletions of the screen, the outcome of deleting it are in Supplementary Table S1.

### Responses to the environment and to stress

The model writes 28 two-component systems, in which a sensor histidine kinase reads its signal and phosphorylates its response regulator [53, 54]; they are the cell’s means of answering a change in its surroundings. In version 1.0 two are written from measured constants: EnvZ/OmpR, which reads osmolarity and sets the balance of the OmpF and OmpC porins (Listing 4), and PhoR/PhoB, which reads phosphate and controls the Pho regulon. Twenty-four more, NtrB/NtrC among them, are generated from one template whose constants obey a stated rule, and two, KdpD/KdpE and CheA/CheY, keep an older template. In EnvZ/OmpR and PhoR/PhoB, several constants were set from the responses they are later compared with (Table 6). The EnvZ kinase and its osmolality factor were set to give the measured OmpR*∼*P fractions of 5% and 27% [46]; the EnvZ phosphatase runs at about 67 times its rate *in vitro*, so that OmpR*∼*P reaches its maximum about 2 min after the stimulus, as measured [46]; and the *ompC* activation and *ompF* repression constants were set to the 3.4-fold rise and the halving of their expression measured by Jovanovich *et al.* [47]. The PhoR kinase constant is the one Gao and Stock fitted to PhoB phosphorylation *in vivo* [48], and the Pho promoter affinities come from the same studies, with three model values chosen to keep the measured order [49, 48]. These tests therefore check that the whole cell keeps the responses written into the modules. EnvZ is also the phosphatase of OmpR*∼*P, and this architecture keeps the OmpR*∼*P concentration nearly independent of the levels of OmpR and EnvZ while EnvZ is much scarcer than OmpR, as in the cell [55, 56]: in a test of the module on its own, OmpR*∼*P stays within 2.2% over a fourfold range of OmpR and an eightfold range of EnvZ.

**Table 6.** Responses of two-component systems to a change in the medium at *t* = 3,000 s in the whole cell, against measurement, on development versions D (EnvZ/OmpR) and E (PhoR/PhoB), built before version 1.0. Constants of both modules were set from these measurements (text), and where the model and the measurement report different quantities, the table names each.

| Change (system) | Model | Measured |
|---|---|---|
| NaCl 50 to 350 mM (EnvZ/OmpR) | OmpR∼P 5.5% to 34%, half-rise 60 s; *ompC* mRNA 2.0-fold, *ompF* mRNA 0.46-fold | OmpR∼P 5% and 27% (20% sucrose, taken as 0.3 M NaCl), at its maximum about 2 min after procaine [46]; *ompC-lac* expression rate 3.4-fold until about 40 min, then 2-fold, and *ompF-lac* expression halved [47] |
| Phosphate to 1 *µ*M (PhoR/PhoB) | PhoB∼P 2.8 *µ*M at 20–50 min, 4.2 *µ*M by 83 min; PstS and UgpB about 20 *µ*M; *phnC* last | total PhoB half-rise about 25 min [48]; promoter order with *phnC* last [49] |

On development version D, when the NaCl of the medium rises from 50 to 350 mM at *t* = 3,000 s, OmpR*∼*P in the whole cell goes from 5.5% to 34% of OmpR, with half of the rise in 60 s, against measured fractions of 5% in low-osmolarity medium and 27% after 20% sucrose is added, which the model takes as the same osmotic step as 0.3 M NaCl, and a rise after procaine treatment that reaches its maximum in about 2 min and then holds [46]. The *ompC* transcript doubles and the *ompF* transcript falls to 0.46 of its level, against a 3.4-fold rise in the expression rate of an *ompC*-*lac* fusion in the 40 min after 0.3 M NaCl is added, which then settles at twofold, and a twofold fall in *ompF*-*lac* expression [47]; the model reports transcript levels, and the measurements are expression rates of fusions. Growth and ATP do not change. The step acts through EnvZ/OmpR alone: the model has no osmotic water flow, potassium uptake or growth arrest yet, so the comparison is made in a cell whose physiology differs from that of cells in their growth lag after the step.

On development version E, when the phosphate of the medium falls to 1 *µ*M, periplasmic phosphate follows within 60 s, PhoB*∼*P reaches 2.8 *µ*M 20 to 50 min later and 4.2 *µ*M by 83 min, the periplasmic binding proteins PstS and UgpB of their ABC transporters rise from tens of nanomolar to about 20 *µ*M, near the starved levels their constants were set to give, and the Pho promoters turn on in the measured order, *phnC* last [49], an order set by the promoter affinities. Measured total PhoB rises with a half-time of about 25 min after phosphate runs out [48]; the model’s figure is the PhoB*∼*P concentration, and its half-rise time of total PhoB was not recorded. PhoR reads periplasmic phosphate directly, without the PstSCAB transporter and PhoU through which PhoR senses phosphate in the cell. Cytoplasmic phosphate stays clamped in this test, which therefore covers signaling and expression only, and it ran in the second fast-species setting (Methods), in which the unperturbed cell’s ATP also stays below its measured level.

On a development version after 1.0, NtrB/NtrC, which reads the cell’s carbon and nitrogen status through 2-oxoglutarate and glutamine and controls glutamine synthetase, was rebuilt on measured values. In the rich medium, lowering ammonium does not limit the cell’s nitrogen, which comes from the medium’s amino acids; with 2-oxoglutarate imposed at 1.5 mM, near the 1.4 mM measured in nitrogen-limited cells [57], and glutamine at 0.48 mM, a model value, NtrC*∼*P rises from 2.2 to 27 nM within 3 min, and glutamine synthetase reaches 8.7 *µ*M within an hour, about three times its level in the rich medium. This test has no measured output to compare with yet.

The 24 generated systems follow one template whose constants obey a stated rule. In the rich medium their regulators sit 4 to 8% phosphorylated, but their outputs can rise at most 1.5-fold, and several read a stand-in signal or one the cell does not hold: among them, the envelope-stress sensors CpxA and RcsC and the acid sensor EvgS read the clamped H^+^, and the copper, zinc, nitrate, nitrite, trimethylamine *N*-oxide, hydrogen peroxide and ferrous iron sensors read signals the model does not hold. These systems do not yet answer their stimuli. CheA/CheY, on the older template, is a placeholder that departs from the cellular pathway in sign and in output. In the model, cytoplasmic aspartate must bind CheA before CheA autophosphorylates, so aspartate activates the kinase, and CheY*∼*P binds the *fliM* gene and recruits *σ* ^70^ as a transcriptional activator would. In the cell, attractants such as aspartate, sensed by the transmembrane receptor Tar, inhibit CheA, CheZ dephosphorylates CheY*∼*P [58], and CheY*∼*P binds the switch protein FliM [59] at the flagellar motor to reverse its rotation [60, 61]. The recruitment step leads to no transcription in version 1.0, since no reaction uses its product, and *fliM* is transcribed by the *σ* ^70^ holoenzyme like most genes. With the *fliA* promoter set to zero (above), the flagellar sigma factor is not made either, and the model has no flagellar rotation, so version 1.0 has no chemotactic response. Oxygen is the largest gap: removing it drops ATP to near zero within minutes, since the cell’s fermentative enzymes are present only as placeholders and carry no appreciable flux, and holding it at 0.5 or 5 *µ*M stalls the simulation, so the response of ArcB/ArcA, which reads the redox state of the quinone pool, cannot yet be tested in the whole cell.

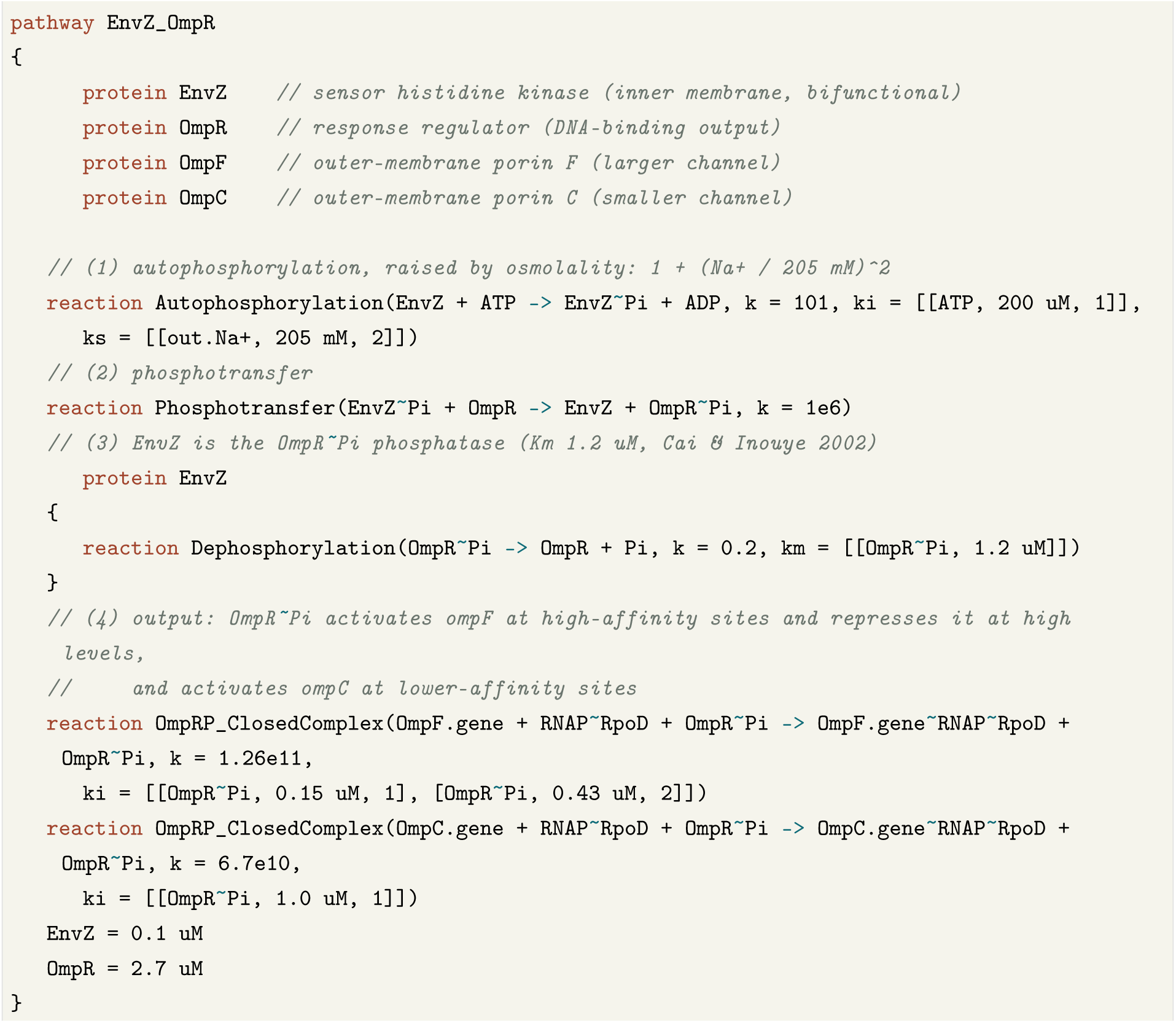

**Listing 4.** Sensing osmolarity: the EnvZ/OmpR two-component system of version 1.0. The medium’s Na^+^ stands for osmolality and raises EnvZ autophosphorylation through a ks term on out.Na+; EnvZ transfers the phosphate to OmpR and is also the OmpR*∼*P phosphatase; OmpR*∼*P sets *ompF* and *ompC* transcription through ki terms at its binding affinities. Code lines verbatim from the model’s EnvZ/OmpR module; comments shortened, and the connector row, the spontaneous hydrolysis of OmpR*∼*P, the promoter rows of *envZ* and *ompR* and the switched-off default promoter rows left out. The phosphotransfer constant is a model value about a hundredth of the measured one, lowered to stay within one simulation tick (Methods); the phosphatase rate and the two output constants are model values set from the responses of Table 6.

### Perturbations written as edits

Because the whole system is code, a perturbation that takes days at the bench is an edit. This section shows how such experiments are written, and it reports no result from them. Thymidylate synthase (ThyA) supplies dTMP for DNA replication. Following the CRISPRi epistasis design of Otto *et al.* [62], its expression is titrated by scaling the holoenzyme-docking constant at its promoter, or the gene is deleted outright (Listing 5); the enzyme itself is declared once. The envelope and drug-target subsystems carry placeholder antibiotic routes, whose response this paper does not evaluate: a lumped penicillin-binding protein binds imipenem reversibly (PBP∼Imipenem), so that wall synthesis stops, and gentamicin enters through a passive channel and binds the ribosome.

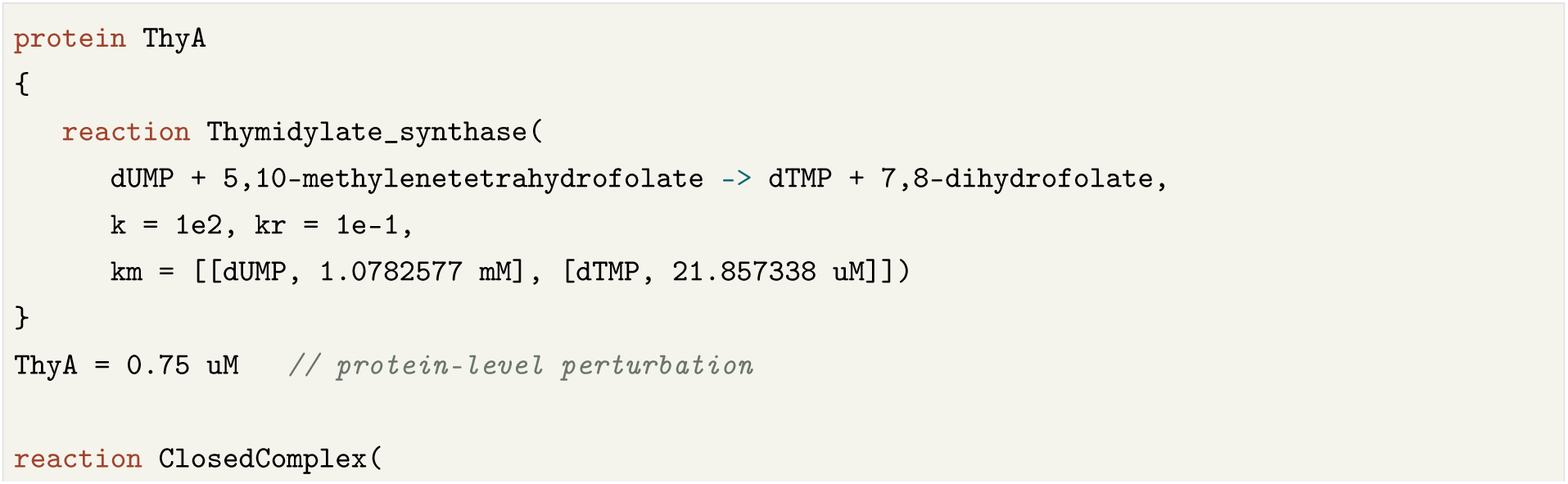
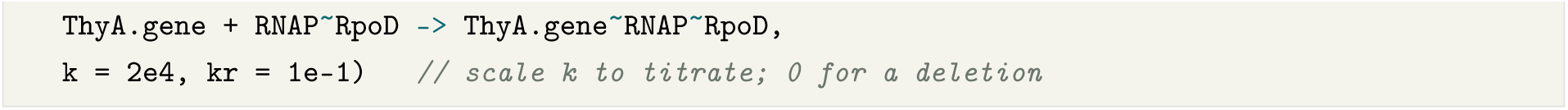

**Listing 5.** A perturbable module. ThyA’s kinetics are declared once, with model values for its constants that cite no source, and the reaction is written reversible although the step runs one way in the cell; the gene is retuned or deleted by editing a single line. From the model’s pyrimidine metabolism module, commented-out alternatives left out; the last two comments are ours.

## Discussion

Writing *E. coli* in a language whose nouns are biological makes the organism itself the deliverable. The model is readable, since an enzyme is an enzyme and a gene expresses through explicit reactions; composable, since its subsystems combine by containment over one shared molecular pool; and reproducible, since a run repeats from its seed and one declaration ties a gene’s expression to its catalysis, so the two cannot drift apart. Most of the agreement with measurement reported here was put in by calibration, starting values, held pools or gates written from the data (Table 4), so it shows that the whole cell keeps what its parts were set to do. The tests that come next are predictions against data the model was not set to: genome-wide deletions scored for essential and non-essential genes alike, other media, and the growth laws that relate the composition of a cell to its growth rate.

### What the model does not yet do

The model reports where it departs from measurement. Its largest limitation is its clamps: 20 of the 31 pools held at fixed values in version 1.0 stand in for synthesis not yet written, among them coenzyme A, the folates and the pentose phosphate intermediates. A clamped pool agrees with measurement by construction, so that agreement is not a result, and a clamp can feed or drain the reactions around it unseen until it is released, as the carbamoyl phosphate and coenzyme A clamps did. Ten of the measured pools fall as fast as growth dilutes them, because the model makes them at almost no rate. Version 1.0 does not run the pentose phosphate pathway, writes its glucose phosphotransferase system only in part, and has no pH: H^+^ is a held supply pool. The cell replicates one round at a time from an origin held loaded, at a fork rate set to fit its cycle, with no DnaA control of initiation and no D period, and it divides at a fixed length. The cell does not yet degrade its ribosomes when its growth arrests. The model has no low-oxygen state, since its fermentative enzymes are only placeholders, and no osmotic water flow. Of its two-component systems only EnvZ/OmpR and PhoR/PhoB are built on measured constants, and they were tested on development versions with constants set from the responses they reproduce. DNA repair, toxin activity, chemotaxis and the prophages are placeholders, and CheA/CheY carries errors of sign and output. NADP^+^ runs about 620 times above its measured pool and NADH about five times, and eight pools are held at ratios that Michaelis constants chosen at raised starting values impose. Many rate constants are model values or placeholders (Methods) and await systematic training against condition-resolved omics, and several subsystems capture the initiating mechanism without its full downstream structure. Every 50S subunit consumes the zinc-regulated paralogs YkgM and RpmJ2 beside L31 and L36, and LysU charges no tRNA. Proteins counted with their complexes run above the LB proteome, with ribosomal proteins high and the most abundant other proteins low, and the cell is smaller than an LB-grown cell. Stochastic trajectories need ensembles for population read-outs such as kill curves, and the numerical method has not been tested for convergence across tick sizes (Methods).

### Version 2 and a shared model

The next version closes these gaps in order: release the pools still held to their own synthesis, bring the metabolic core onto measured kinetics pathway by pathway, rerun the genome-wide deletion screen on the current model and score it for essential and non-essential genes alike, and extend validation from one rich medium to growth, flux and expression across conditions, the microaerobic state among them. No single group commands the domain expertise, the measurements and the experimental validation that a whole cell needs. Written in a shared language, the model lets scientists combine their expertise, experiments and data, each contribution tested against the rest of the cell before it joins. The companion paper describes the shared library of organism code, with the virtual *E. coli* as a baseline that others can extend and benchmark and a mechanistic model of the human as its long-term goal [25]; we plan to extend the library to further microbes and industrial strains and to eukaryotic cells. AI-based efforts pursue virtual cells by learning from data [63].

## Methods

### Sources

The genome is the K-12 MG1655 reference (GenBank U00096.3) [1, 26]; each gene’s nucleotides and residues (Listing 2) are counted from the coding and protein sequences of the same record, matched to the model’s genes by UniProt accession or gene name. Primary sigma factors follow RegulonDB [7], with a gene that has both filed under its alternative sigma by convention (Table 3); protein abundances follow PaxDb [40] and the proteome of Schmidt *et al.* [15]; kinetic constants come from BRENDA [10] and SABIO-RK [11] and the primary literature cited in the model files; metabolite concentrations follow Bennett *et al.* [14]; and essential-gene calls follow Goodall *et al.* [45]. Where a value has a source, the model file names it. Of the 41,649 reactions of version 1.0, 35,463 are per-gene central-dogma rows (promoter binding, open complex, elongation, polymerase release, mRNA decay, ribosome binding, protein synthesis and protein decay) and 6,186 lie outside them. We traced 6,176 of these 6,186 to the reaction statement that defines them and sorted each into one category by what its code comments say about its values. Of these, 32 cite a source for all their values, 94 cite a source for some values and mark the others as model values, and 119 are marked as model values with no citation. The other 5,931 name no source: 1,653 of them have a forward rate constant of zero, 2,384 use the shared default rate constant *k* = 0.01 with no Michaelis constant, and 1,894 have other values. A comment counts as citing a source when it gives an author and year, the words *et al.*, a DOI or a PMID, or names BRENDA, SABIO- RK, BioNumbers or EcoCyc; tags that only say where the equation of a reaction came from (a UniProt annotation, an EC number, a BiGG identifier) do not count. In all, 249 reactions are tagged in the code as proposed by a language model; none cites a source, 131 use the default rate constant and 118 other values. A further 391 carry a template-placeholder tag (186 from EC-number templates, 106 from role templates and 99 from mobile-element templates). Of the reactions with a zero rate constant, 816 are rows of the transcription-factor module, about half of its rows, so about half of the regulation written there is switched off in version 1.0. The counting script is in Supplementary File 1.

### Writing each gene’s central dogma

The four files that write out each gene’s central dogma (Listing 1) are generated by scripts and kept as L++ code. The cell is first compiled with the sigma-class templates of transcription and translation, and every row those templates expand for a gene is written out as an explicit row of that gene, except a step that a module writes for the gene itself. The class declarations of the modules are rewritten as plain protein declarations, so that a gene of an alternative sigma factor keeps its own promoter rows, and the template reactions are commented out. Each row’s generic composition is then replaced by the gene’s own nucleotide and residue counts (Sources). A check of the compiled cell confirms, gene by gene, that transcription and decay, and synthesis and degradation, return the nucleotides and amino acids they take. The eight scripts are in Supplementary File 1. Run by the same scripts on a later development version, the four files carry the same rows as in version 1.0 for every gene but 27 rows of 12 genes, which modules of that version write themselves.

### Simulation

Whole-cell runs of *E. coli* version 1.0 compile, with lppc, an experiment that places one cell in a dish of the defined rich medium with 4.1 mM glucose, and run it with lpps at resolution level 6 (-r 6), printing every 100 s (-i 100), to *t* = 20,000 s or longer, with seeds 1 and 2. Compiling the cell takes about 50 s on one core. lppc itself peaks at about 4.6 GB of memory; the C++ build of the generated simulation takes about 6 s of that time and 0.3 GB. Free and charged tRNAs and a set of central metabolites are advanced as fast species (the environment variable LPPS_FAST_SPECIES), and LPPS_VC_KM=1 lets the km of a countflux reaction saturate it in its substrates, as transcript and protein elongation require. This is the standard setting; a second setting, used earlier in development, also advances the folates as fast species and lets lpps add the fast species it detects (LPPS_AUTO_FAST=0.2). The dividing population is followed as a count of identical cells. Supplementary File 1 gives the commands step by step, with the analysis scripts.

### Medium

The medium is a modified form of MOPS EZ Rich defined medium (Teknova M2105), itself a modification of the medium of Neidhardt *et al.* [50]. It holds the recipe’s 19 amino acids at its concentrations (serine 10 mM, arginine 5.2 mM, the others 0.1 to 0.8 mM), adenine, cytosine, guanine and uracil at 0.2 mM each, and Na^+^ 50 mM, NH^+^ 9.5 mM, phosphate 1.32 mM, Mg^2+^ 0.53 mM, sulfate 0.29 mM and Ca^2+^ 0.5 *µ*M, as in the recipe. Glucose is 4.1 mM and glutamate 11 mM, against 11.1 mM (0.2%) and 0.6 mM in the recipe. The vitamins are nicotinamide (98 *µ*M), pantothenate (49 *µ*M), pyridoxine (17 *µ*M), thiamine (15 *µ*M), lipoate (9.7 *µ*M), riboflavin (2.1 *µ*M), biotin (0.49 *µ*M) and folate (0.18 *µ*M); the recipe has 10 *µ*M each of thiamine, calcium pantothenate, *p*-aminobenzoate, *p*-hydroxybenzoate and 2,3-dihydroxybenzoate, and the last three are absent from the model medium. Iron is 10 nM against 10 *µ*M

FeSO_4_ in the recipe, and the MOPS and tricine buffers are not represented. H^+^, O_2_ and CO_2_ are held at 1.2 mM in the medium, the periplasm and the cytoplasm for the whole run (Clamped pools and their release). The dish holds 100 mL, and at *t* = 20,000 s every medium value is unchanged at the precision the run prints.

### Rate laws and the numerical method

A reaction with Michaelis constants runs on the species its km list names alone, each entering as its concentration over its Michaelis constant. Its forward and backward terms share one denominator, so a reversible reaction comes to rest where *k* ∏*_s_ c_s_/K_s_* = *k_r_* ∏*_p_ c_p_/K_p_*, and reactants and products outside the list, such as ATP, H^+^ or carbamoyl phosphate where they are not listed, do not enter its rate. Lpps advances every reaction in fixed ticks of 2*^−^*^6^ s at resolution level 6. In a tick, counted species fire whole events drawn from the run’s seeded generator; when a draw would take a counted species below zero by half a molecule or more, the tick is drawn again, up to five times, and after that a depletion guard scales the reactions that consume it. In each of the two 20,000 s runs of version 1.0 (1,280,001 ticks), the guard acted between 800,001 and 900,000 times; lpps logs one action in 100,000, and each logged action scaled one to three species, except the first in seed 2, which scaled eleven at *t* = 0. lpps also estimates the tick each reaction would need so that no amount changes too much in one tick; for this model it asks for level 14 (2*^−^*^14^ s) at the start, and later for level 408 when an amount approaches zero. The runs keep level 6, and the redraws and the guard absorb the difference; each finer level halves the tick and doubles the number of ticks, and no convergence test across levels has been run on version 1.0. At least one rate constant was lowered for the tick: EnvZ transfers its phosphate to OmpR at 10^6^ M*^−^*^1^ s*^−^*^1^, against about 10^8^ M*^−^*^1^ s*^−^*^1^ measured, so that the step stays within one tick (Listing 4).

### Model versions

Results from development versions built before version 1.0 carry neutral labels: A, the version of Figure 3 and of the representative deletions; B, the version of the screen of 189 deletions; C, the version of the cell volume; D and E, the versions of the NaCl and phosphate steps. A rebuilt NtrB/NtrC module ran on a development version after 1.0. Version 1.0 writes each gene’s central dogma out (Listing 2); the routes of Table 5 and the census of the sources of its values are counted on it.

### Analysis

Divisions are read from the cell count in the run’s output. Pools are averaged over the stated window of the cytoplasmic compartment and compared with Bennett *et al.* [14] as log_2_ fold changes; proteins, each counted with the complexes it belongs to, are compared with the measured proteome. Negativeconcentration events are those lpps counts and reports at the end of a run. A deletion sets the gene’s transcription initiation to zero (lpps --knockout) and removes the protein already present by a degradation event at *t* = 0. Each deletion is scored by its cell count at the end of the run against the wild type at the same time: a stop is one cell, a slowdown fewer cells than the wild type, and growth the wild-type count. The 189 deletions of the screen on version B ran on seed 1 to *t* = 1,500 s (64 deletions; wild type 2 cells), 3,000 s (101; 4 cells) or 5,000 s (24; 16 cells), and the representative deletions on version A ran on seed 1 to *t* = 3,000 s.

### Releasing a clamp and testing a unit

A pool is released by writing the pathway that makes it, with measured kinetics where they exist, removing its fixed value, giving each reaction that consumes it a Michaelis constant for it, and checking the equilibrium that its rate constants imply. A release, like each unit that brings a pathway to measured values, is tested in the whole cell before it joins the model: seeds 1 and 2 to *t* = 6,000 s against the unchanged model, the deletions that touch the pathway, and no amount driven below zero, in both fast-species settings when the change touches purine or folate metabolism. The pair of 20,000 s runs and the comparisons of pools and proteins then run on the combined build.

### Use of AI tools

The authors used Claude (Anthropic), through Claude Code, to draft and revise the text, to write and revise parts of the model code under the authors’ direction, and to run and analyze the simulations. The authors directed this work, checked every reference and every numerical result reported in the paper against its source, and take responsibility for the paper. The model’s parameters carry a source note where they have one, and the census in Methods gives how many do, including the 249 reactions whose equations and values a language model inferred, which the code marks as such. An AI tool is not an author of this work.

## Data availability

The model’s L++ code, *E. coli* version 1.0, is in Supplementary File 1, with the data files lpps reads, the scripts that write each gene’s central dogma, the analysis scripts and the script that counts the sources of the model’s values, and a guide that reproduces step by step the whole-cell runs of version 1.0 and the results drawn from them: the division times, the metabolite and protein comparisons of Table 4 and Figure 2, and Supplementary Tables S2 and S3. The code is released under the Life Code License of Omphalos Lifesciences Inc, whose text Supplementary File 1 includes. Results from development versions A to E (Figure 3, the deletion screen and the representative deletions, the cell volume of version C and the responses of Table 6) come from runs of those versions, which Supplementary File 1 does not reproduce; it carries the data plotted in Figure 3 and the outcome of each deletion of the screen. Supplementary File 1 also carries lppc and lpps 1.0.0 for Linux on 64-bit ARM, the build released with the companion paper [25]; builds for Linux on x86-64 and for Windows are available from the corresponding authors on request. Supplementary Tables S1 to S3 give the 377 essential-gene candidates, the 95 measured metabolite pools and the 31 pools still held at fixed values; Supplementary File 1 carries them as one page and as tab-separated files.

## Acknowledgements

This research was supported by the Bio & Medical Technology Development Program of the National Research Foundation (NRF) and funded by the Korean government (MSIT) (No. RS-2022-NR067509).

## Author contributions

D.M.L. and D.K. built the model. All authors analyzed the results and prepared the manuscript.

## Supplementary material

**Supplementary File 1.** The L++ code of *E. coli* version 1.0 with lppc and lpps for Linux on 64-bit ARM, and how to reproduce the whole-cell runs and analyses of version 1.0: the commands, the scripts and the data behind the figures.

**Supplementary Table S1.** The 377 essential-gene candidates, with each screen’s call, the route by which growth depends on each and the outcome of deleting it on development version B.

**Supplementary Table S2.** The 95 measured metabolite pools against the model.

**Supplementary Table S3.** The 31 pools still held at fixed values.

