## Supplementary File 1 for "A Genome-Scale Virtual *Escherichia coli* in the L++ Language": Supplementary_Tables_S1-S3.html

Supplementary Tables S1 to S3: A Genome-Scale Virtual Escherichia coli in the L++ Language 

Supplementary Tables S1 to S3 of "A Genome-Scale Virtual *Escherichia coli* in the L++ Language". The same tables are in this folder as tab-separated files.

### Supplementary tables

#### Supplementary Table S1. The 377 essential-gene candidates

Every candidate that any of the Keio, PEC and TraDIS screens calls essential (Goodall *et al.* 2018): 248 in
three screens, 45 in two, 46 in one and 38 in none. For each, the screens that call it, the route by which the model's
growth depends on it as counted on development version B, and the outcome of deleting it on that version (a dash
where the deletion was not run). Version 1.0's counts are those of Table 5: there the cell-wall genes were
rewired onto the peptidoglycan pathway, and *polA*, *priA*, *recA* and *rnhA* have no route.
Eleven deletions that the screen's record lists as slowing end at the wild type's cell count, and their rows say so.

| Gene | Protein | Screens | Route to growth | Deletion (earlier version) |
| --- | --- | --- | --- | --- |
| **Aminoacyl-tRNA Synthetases** (22) | | | | |
| *alaS* | Alanyl-tRNA Synthetase | PEC, TraDIS | sole tRNA charging enzyme | – |
| *argS* | Arginyl-tRNA Synthetase | Keio, PEC, TraDIS | sole tRNA charging enzyme | – |
| *asnS* | Asparaginyl-tRNA Synthetase | Keio, PEC, TraDIS | sole tRNA charging enzyme | – |
| *aspS* | Aspartyl-tRNA Synthetase | Keio, PEC, TraDIS | sole tRNA charging enzyme | – |
| *cysS* | Cysteinyl-tRNA Synthetase | Keio, PEC, TraDIS | sole tRNA charging enzyme | – |
| *glnS* | Glutaminyl-tRNA Synthetase | Keio, PEC, TraDIS | sole tRNA charging enzyme | – |
| *gltX* | Glutamyl-tRNA Synthetase | Keio, PEC, TraDIS | sole tRNA charging enzyme | – |
| *glyQ* | Glycyl-tRNA Synthetase Alpha Subunit | Keio, PEC, TraDIS | sole tRNA charging enzyme | – |
| *glyS* | Glycyl-tRNA Synthetase Beta Subunit | PEC, TraDIS | presence gate | stops |
| *hisS* | Histidyl-tRNA Synthetase | Keio, PEC, TraDIS | sole tRNA charging enzyme | – |
| *ileS* | Isoleucyl-tRNA Synthetase | PEC, TraDIS | sole tRNA charging enzyme | – |
| *leuS* | Leucyl-tRNA Synthetase | Keio, PEC, TraDIS | sole tRNA charging enzyme | stops |
| *lysS* | Lysyl-tRNA Synthetase | TraDIS | sole tRNA charging enzyme in the model (LysU has no charging reaction, a model gap) | – |
| *metG* | Methionyl-tRNA Synthetase | Keio, PEC, TraDIS | sole tRNA charging enzyme | – |
| *pheS* | Phenylalanyl-tRNA Synthetase Alpha Subunit | Keio, PEC, TraDIS | sole tRNA charging enzyme | – |
| *pheT* | Phenylalanyl-tRNA Synthetase Beta Subunit | Keio, PEC, TraDIS | presence gate | stops |
| *proS* | Prolyl-tRNA Synthetase | Keio, PEC, TraDIS | sole tRNA charging enzyme | – |
| *serS* | Seryl-tRNA Synthetase | Keio, PEC, TraDIS | sole tRNA charging enzyme | – |
| *thrS* | Threonyl-tRNA Synthetase | Keio, PEC, TraDIS | sole tRNA charging enzyme | – |
| *trpS* | Tryptophanyl-tRNA Synthetase | Keio, PEC, TraDIS | sole tRNA charging enzyme | – |
| *tyrS* | Tyrosyl-tRNA Synthetase | Keio, PEC, TraDIS | sole tRNA charging enzyme | – |
| *valS* | Valyl-tRNA Synthetase | Keio, PEC, TraDIS | sole tRNA charging enzyme | – |
| **Aromatic Amino Acid Biosynthesis (Shikimate Pathway)** (5) | | | | |
| *aroA* | EPSP Synthase | (none) | reactions | slows |
| *aroB* | 3-Dehydroquinate Synthase | (none) | reactions | slows |
| *aroC* | Chorismate Synthase | (none) | reactions | slows |
| *aroE* | Shikimate Dehydrogenase | (none) | reactions | slows |
| *aroK* | Shikimate Kinase | (none) | reactions | slows as recorded; ends at the wild-type count |
| **Cell Division** (12) | | | | |
| *ftsA* | Z-Ring Membrane Anchor | Keio, PEC, TraDIS | presence gate | stops |
| *ftsB* | Divisome Protein FtsB | Keio, PEC, TraDIS | presence gate | – |
| *ftsE* | Divisome ATPase FtsE | Keio, PEC | presence gate | – |
| *ftsI* | Penicillin-Binding Protein 3 (PBP3) | Keio, PEC, TraDIS | presence gate | stops |
| *ftsK* | DNA Translocase / Divisome Component | Keio, PEC | presence gate | stops |
| *ftsL* | Divisome Protein FtsL | Keio, PEC, TraDIS | presence gate | – |
| *ftsN* | Late Divisome Trigger | Keio, PEC | presence gate | stops |
| *ftsQ* | Divisome Assembly Scaffold | Keio, PEC, TraDIS | presence gate | stops |
| *ftsW* | Septal Lipid II Flippase/Polymerase | Keio, PEC, TraDIS | presence gate | stops |
| *ftsX* | Divisome Membrane Protein FtsX | Keio, PEC | presence gate | – |
| *ftsZ* | Z-Ring GTPase (Tubulin Homolog) | Keio, PEC, TraDIS | presence gate | stops |
| *zipA* | Z-Ring Membrane Anchor | Keio, PEC, TraDIS | presence gate | stops |
| **Central Metabolism (Chokepoint Enzymes)** (15) | | | | |
| *aceF* | Pyruvate Dehydrogenase E2 Subunit | TraDIS | reactions | slows |
| *can* | Carbonic Anhydrase | Keio, PEC, TraDIS | presence gate | – |
| *eno* | Enolase | Keio, PEC, TraDIS | reactions | grows |
| *fbaA* | Fructose-Bisphosphate Aldolase | Keio, PEC, TraDIS | reactions | slows as recorded; ends at the wild-type count |
| *gapA* | Glyceraldehyde-3-Phosphate Dehydrogenase | Keio, PEC, TraDIS | reactions | slows as recorded; ends at the wild-type count |
| *glmM* | Phosphoglucosamine Mutase | PEC, TraDIS | presence gate | – |
| *glmS* | Glutamine-Fructose-6-P Aminotransferase | Keio, PEC, TraDIS | presence gate | – |
| *glmU* | Bifunctional GlcNAc-1-P Uridyltransferase/Acetyltransferase | Keio, PEC, TraDIS | presence gate | – |
| *lpd* | Dihydrolipoamide Dehydrogenase | TraDIS | slowdown | slows as recorded; ends at the wild-type count |
| *pgk* | Phosphoglycerate Kinase | Keio, PEC, TraDIS | reactions | slows |
| *ppa* | Inorganic Pyrophosphatase | Keio, PEC, TraDIS | presence gate | – |
| *rpe* | Ribulose-Phosphate 3-Epimerase | TraDIS | slowdown | – |
| *sucA* | 2-Oxoglutarate Dehydrogenase E1 Subunit | TraDIS | reactions | slows as recorded; ends at the wild-type count |
| *sucB* | 2-Oxoglutarate Dehydrogenase E2 Subunit | TraDIS | reactions | grows |
| *tktA* | Transketolase I | TraDIS | slowdown | slows |
| **Cofactor Biosynthesis** (39) | | | | |
| *birA* | Biotin Ligase and Repressor | Keio, PEC, TraDIS | reactions | grows |
| *coaA* | Pantothenate Kinase | PEC, TraDIS | reactions | grows |
| *coaD* | Phosphopantetheine Adenylyltransferase | Keio, PEC, TraDIS | reactions | grows |
| *coaE* | Dephospho-CoA Kinase | PEC, TraDIS | reactions | grows |
| *dfp* | Phosphopantothenoylcysteine Synthetase/Decarboxylase | Keio, PEC, TraDIS | reactions | grows |
| *folA* | Dihydrofolate Reductase (DHFR) | Keio, PEC, TraDIS | presence gate | stops |
| *folB* | Dihydroneopterin Aldolase | PEC, TraDIS | reactions | grows |
| *folC* | Folylpolyglutamate Synthase | Keio, PEC, TraDIS | reactions | grows |
| *folD* | Methylene-THF Dehydrogenase and Cyclohydrolase | Keio, PEC, TraDIS | reactions | grows |
| *folE* | GTP Cyclohydrolase I | Keio, PEC, TraDIS | reactions | grows |
| *folK* | 2-Amino-4-Hydroxy-6-Hydroxymethyldihydropteridine Pyrophosphokinase | Keio, PEC | reactions | grows |
| *folP* | Dihydropteroate Synthase | TraDIS | reactions | grows |
| *hemA* | Glutamyl-tRNA Reductase | Keio, PEC, TraDIS | presence gate | stops |
| *hemB* | Delta-Aminolevulinic Acid Dehydratase | Keio, PEC, TraDIS | presence gate | – |
| *hemC* | Porphobilinogen Deaminase | Keio, PEC, TraDIS | presence gate | – |
| *hemD* | Uroporphyrinogen III Synthase | Keio, PEC, TraDIS | presence gate | – |
| *hemE* | Uroporphyrinogen Decarboxylase | TraDIS | slowdown | slows |
| *hemG* | Protoporphyrinogen Oxidase | Keio, PEC, TraDIS | presence gate | – |
| *hemH* | Ferrochelatase | Keio, PEC, TraDIS | presence gate | – |
| *hemL* | Glutamate-1-Semialdehyde Aminotransferase | Keio, PEC, TraDIS | presence gate | – |
| *metK* | S-Adenosylmethionine Synthetase | Keio, PEC, TraDIS | presence gate | stops |
| *nadD* | Nicotinate-Nucleotide Adenylyltransferase | Keio, PEC, TraDIS | reactions | grows |
| *nadE* | NAD Synthetase | Keio, PEC, TraDIS | reactions | grows |
| *nadK* | NAD Kinase | Keio, PEC, TraDIS | reactions | grows |
| *pdxH* | Pyridoxine 5'-Phosphate Oxidase | TraDIS | slowdown | – |
| *ribA* | GTP Cyclohydrolase II | Keio, PEC, TraDIS | reactions | grows |
| *ribB* | 3,4-Dihydroxy-2-Butanone-4-Phosphate Synthase | Keio, PEC | reactions | grows |
| *ribC* | Riboflavin Synthase | Keio, PEC, TraDIS | reactions | grows |
| *ribD* | Riboflavin Deaminase and Reductase | Keio, PEC, TraDIS | reactions | grows |
| *ribE* | Lumazine Synthase | Keio, PEC, TraDIS | reactions | slows as recorded; ends at the wild-type count |
| *ribF* | Riboflavin Kinase / FAD Synthetase | Keio, PEC, TraDIS | reactions | grows |
| *thiL* | Thiamine-Phosphate Kinase | Keio, PEC, TraDIS | reactions | grows |
| *ubiA* | 4-Hydroxybenzoate Octaprenyltransferase | Keio, PEC, TraDIS | presence gate | stops |
| *ubiB* | Ubiquinone Synthesis Protein UbiB | Keio, TraDIS | presence gate | – |
| *ubiD* | Octaprenylhydroxybenzoate Decarboxylase | Keio, TraDIS | presence gate | – |
| *ubiE* | Ubiquinone/Menaquinone Methyltransferase | TraDIS | slowdown | – |
| *ubiG* | Ubiquinone Biosynthesis Methyltransferase | TraDIS | slowdown | – |
| *ubiH* | 2-Octaprenyl-6-Methoxyphenol Hydroxylase | TraDIS | slowdown | – |
| *ubiX* | Flavin Prenyltransferase | TraDIS | slowdown | – |
| **DNA Replication & Chromosome Maintenance** (18) | | | | |
| *dnaA* | Chromosomal Replication Initiator | Keio, PEC, TraDIS | presence gate | stops |
| *dnaB* | Replicative Helicase | Keio, PEC, TraDIS | presence gate | stops |
| *dnaC* | Helicase Loader | Keio, PEC, TraDIS | presence gate | stops |
| *dnaE* | DNA Polymerase III Alpha Subunit | Keio, PEC, TraDIS | presence gate | stops |
| *dnaG* | DNA Primase | PEC, TraDIS | presence gate | stops |
| *dnaN* | Beta Sliding Clamp | Keio, PEC, TraDIS | presence gate | stops |
| *dnaT* | Primosome Protein DnaT | PEC, TraDIS | presence gate | – |
| *dnaX* | Clamp Loader (Tau/Gamma) | Keio, PEC, TraDIS | presence gate | stops |
| *hda* | RIDA regulator | PEC, TraDIS | presence gate | stops |
| *holA* | Clamp Loader Delta Subunit | Keio, PEC, TraDIS | presence gate | – |
| *holB* | Clamp Loader Delta-Prime Subunit | Keio, PEC, TraDIS | presence gate | stops |
| *lexA* | SOS Repressor | Keio, PEC, TraDIS | reactions | grows |
| *ligA* | DNA Ligase (NAD-dependent) | Keio, PEC, TraDIS | presence gate | stops |
| *polA* | DNA Polymerase I | PEC | slowdown (no route in version 1.0) | slows |
| *priA* | Primosomal Protein N' | PEC | slowdown (no route in version 1.0) | slows |
| *recA* | Recombinase / SOS Regulator | (none) | slowdown (no route in version 1.0) | slows |
| *rnhA* | RNase HI | (none) | slowdown (no route in version 1.0) | slows |
| *ssb* | Single-Stranded DNA-Binding Protein | Keio, PEC, TraDIS | presence gate | stops |
| **DNA Topology** (5) | | | | |
| *gyrA* | DNA Gyrase Subunit A | Keio, PEC, TraDIS | presence gate | stops |
| *gyrB* | DNA Gyrase Subunit B | Keio, PEC, TraDIS | presence gate | stops |
| *parC* | Topoisomerase IV Subunit A | PEC, TraDIS | presence gate | stops |
| *parE* | Topoisomerase IV Subunit B | Keio, PEC, TraDIS | presence gate | stops |
| *topA* | Topoisomerase I | Keio, PEC, TraDIS | presence gate | – |
| **Diaminopimelate & Lysine Precursor Pathway** (6) | | | | |
| *asd* | Aspartate-Semialdehyde Dehydrogenase | Keio, PEC, TraDIS | reactions | slows |
| *dapA* | 4-Hydroxy-Tetrahydrodipicolinate Synthase | Keio, PEC, TraDIS | presence gate | stops |
| *dapB* | 4-Hydroxy-Tetrahydrodipicolinate Reductase | Keio, PEC, TraDIS | presence gate | stops |
| *dapD* | Tetrahydrodipicolinate Succinylase | Keio, PEC, TraDIS | presence gate | stops |
| *dapE* | N-Succinyl-Diaminopimelate Desuccinylase | Keio, PEC, TraDIS | presence gate | stops |
| *dapF* | Diaminopimelate Epimerase | TraDIS | not wired | grows |
| **Envelope Stress Response & Regulated Proteolysis** (4) | | | | |
| *degS* | Site-1 Regulated Intramembrane Protease | Keio, PEC | presence gate | stops |
| *rpoE* | Sigma-E (Envelope Stress Sigma Factor) | Keio, TraDIS | reactions | grows |
| *rseP* | Site-2 Regulated Intramembrane Protease | Keio, PEC | presence gate | stops |
| *yrfF* | Rcs Pathway Repressor (IgaA) | Keio, PEC, TraDIS | presence gate | – |
| **Iron-Sulfur Cluster Biogenesis** (5) | | | | |
| *erpA* | Fe-S Carrier ErpA | Keio, PEC, TraDIS | presence gate | – |
| *fdx* | Ferredoxin | TraDIS | slowdown | – |
| *hscA* | Fe-S Cluster Chaperone (Hsp70-family) | TraDIS | slowdown | – |
| *iscS* | Cysteine Desulfurase | TraDIS | slowdown | slows |
| *iscU* | Fe-S Cluster Scaffold Protein | TraDIS | slowdown | – |
| **Isoprenoid / Undecaprenyl Carrier Biosynthesis** (12) | | | | |
| *bacA* | Undecaprenyl Pyrophosphate Phosphatase | (none) | not wired | – |
| *dxr* | DXP Reductoisomerase | Keio, PEC, TraDIS | presence gate | – |
| *dxs* | DXP Synthase | Keio, PEC, TraDIS | presence gate | – |
| *fldA* | Flavodoxin 1 | Keio, PEC, TraDIS | reactions | grows |
| *ispA* | Farnesyl Diphosphate Synthase | Keio, PEC, TraDIS | presence gate | – |
| *ispB* | Octaprenyl Diphosphate Synthase | Keio, PEC, TraDIS | presence gate | – |
| *ispD* | MEP Cytidylyltransferase | Keio, PEC, TraDIS | presence gate | – |
| *ispE* | CDP-ME Kinase | Keio, PEC, TraDIS | presence gate | – |
| *ispF* | MEcPP Synthase | Keio, PEC, TraDIS | presence gate | – |
| *ispG* | HMB-PP Synthase | Keio, PEC, TraDIS | presence gate | – |
| *ispH* | HMB-PP Reductase | Keio, PEC, TraDIS | presence gate | – |
| *ispU* | Undecaprenyl Pyrophosphate Synthase | Keio, PEC, TraDIS | presence gate | – |
| **Lipid A / LPS Biosynthesis** (13) | | | | |
| *kdsA* | KDO-8-Phosphate Synthase | Keio, PEC, TraDIS | presence gate | stops |
| *kdsB* | CMP-KDO Synthetase | Keio, PEC, TraDIS | presence gate | stops |
| *kdsC* | KDO-8-Phosphate Phosphatase | PEC | not wired | grows |
| *lapC* | LapC (PbgA/YejM), LPS and Cardiolipin Levels Regulator | Keio, PEC | presence gate | stops |
| *lptF* | LPS Transporter Subunit LptF | Keio, PEC, TraDIS | presence gate | – |
| *lpxA* | UDP-GlcNAc Acyltransferase | Keio, PEC, TraDIS | presence gate | stops |
| *lpxB* | Lipid A Disaccharide Synthase | Keio, PEC, TraDIS | presence gate | stops |
| *lpxC* | UDP-3-O-Acyl-GlcNAc Deacetylase | Keio, PEC, TraDIS | presence gate | stops |
| *lpxD* | UDP-3-O-Acyl-GlcNAc N-Acyltransferase | Keio, PEC, TraDIS | presence gate | stops |
| *lpxH* | UDP-2,3-Diacylglucosamine Hydrolase | Keio, PEC, TraDIS | presence gate | stops |
| *lpxK* | Lipid A 4'-Kinase | Keio, PEC, TraDIS | presence gate | stops |
| *msbA* | Lipid A Flippase | Keio, PEC, TraDIS | presence gate | – |
| *waaA* | KDO Transferase | Keio, PEC, TraDIS | presence gate | stops |
| **Molecular Chaperones & Protein Quality Control** (4) | | | | |
| *ftsH* | Membrane-Bound AAA+ Protease | Keio, PEC, TraDIS | reactions | grows |
| *groL* | GroEL Chaperonin | PEC, TraDIS | presence gate | – |
| *groS* | GroES Co-Chaperonin | Keio, PEC, TraDIS | presence gate | – |
| *grpE* | DnaK Nucleotide Exchange Factor | Keio, PEC, TraDIS | reactions | grows |
| **Nucleoid Structure & Chromosome Segregation** (4) | | | | |
| *ihfA* | Integration Host Factor Alpha Subunit | TraDIS | slowdown | slows |
| *mukB* | SMC-like Chromosome Condensin | Keio, PEC, TraDIS | presence gate | – |
| *mukE* | MukBEF Complex Subunit | Keio, PEC, TraDIS | presence gate | – |
| *mukF* | MukBEF Complex Subunit | Keio, PEC, TraDIS | presence gate | – |
| **Nucleotide Biosynthesis** (30) | | | | |
| *adk* | Adenylate Kinase | Keio, PEC, TraDIS | reactions | stops |
| *carA* | Carbamoyl Phosphate Synthetase (Small Subunit) | (none) | reactions | grows |
| *carB* | Carbamoyl Phosphate Synthetase (Large Subunit) | (none) | reactions | grows |
| *cmk* | Cytidylate Kinase | (none) | reactions | grows |
| *dcd* | dCTP Deaminase | TraDIS | reactions | grows |
| *dut* | dUTP Diphosphatase | Keio, PEC, TraDIS | reactions | stops |
| *gmk* | Guanylate Kinase | Keio, PEC, TraDIS | reactions | stops |
| *guaA* | GMP Synthetase | TraDIS | reactions | grows |
| *ndk* | Nucleoside Diphosphate Kinase | (none) | reactions | stops |
| *nrdA* | Ribonucleotide Reductase (Large Subunit) | Keio, PEC, TraDIS | reactions | stops |
| *nrdB* | Ribonucleotide Reductase (Small Subunit) | Keio, PEC, TraDIS | reactions | stops |
| *prs* | Ribose-Phosphate Diphosphokinase | Keio, PEC, TraDIS | reactions | stops |
| *purB* | Adenylosuccinate Lyase | Keio, TraDIS | reactions | slows |
| *purC* | SAICAR Synthetase | (none) | reactions | grows |
| *purD* | GAR Synthetase | (none) | reactions | grows |
| *purE* | AIR Carboxylase | (none) | reactions | grows |
| *purF* | Amidophosphoribosyltransferase | (none) | reactions | grows |
| *purH* | Bifunctional Purine Biosynthesis Enzyme | (none) | reactions | grows |
| *purK* | N5-CAIR Synthetase | (none) | reactions | grows |
| *purL* | Phosphoribosylformylglycinamidine Synthase | (none) | reactions | grows |
| *purM* | AIR Synthetase | (none) | reactions | grows |
| *purN* | GAR Transformylase | (none) | reactions | grows |
| *pyrB* | Aspartate Carbamoyltransferase | (none) | reactions | slows as recorded; ends at the wild-type count |
| *pyrC* | Dihydroorotase | (none) | reactions | slows |
| *pyrD* | Dihydroorotate Dehydrogenase | (none) | reactions | slows as recorded; ends at the wild-type count |
| *pyrE* | Orotate Phosphoribosyltransferase | (none) | reactions | slows as recorded; ends at the wild-type count |
| *pyrF* | Orotidine-5'-Phosphate Decarboxylase | (none) | reactions | slows as recorded; ends at the wild-type count |
| *pyrG* | CTP Synthetase | Keio, PEC, TraDIS | reactions | stops |
| *pyrH* | UMP Kinase | Keio, PEC, TraDIS | reactions | stops |
| *tmk* | dTMP Kinase | Keio, PEC, TraDIS | reactions | stops |
| **Outer Membrane & Lipoprotein Biogenesis** (17) | | | | |
| *bamA* | Beta-Barrel Assembly Machine Subunit A | Keio, PEC, TraDIS | presence gate | stops |
| *bamD* | BAM Complex Lipoprotein | Keio, PEC, TraDIS | presence gate | stops |
| *lnt* | Apolipoprotein N-Acyltransferase | Keio, PEC, TraDIS | presence gate | – |
| *lolA* | Lipoprotein Chaperone | Keio, PEC, TraDIS | presence gate | stops |
| *lolB* | Outer Membrane Lipoprotein Receptor | Keio, PEC, TraDIS | presence gate | stops |
| *lolC* | Lipoprotein Sorting ABC Transporter | Keio, PEC, TraDIS | presence gate | stops |
| *lolD* | Lipoprotein Export ATPase | Keio, PEC, TraDIS | presence gate | – |
| *lolE* | Lipoprotein Sorting ABC Transporter | Keio, PEC, TraDIS | presence gate | stops |
| *lptA* | LPS Transport Protein A | Keio, PEC, TraDIS | presence gate | stops |
| *lptB* | LPS Transport ATPase | PEC, TraDIS | presence gate | stops |
| *lptC* | LPS Transport Protein C | Keio, PEC | presence gate | stops |
| *lptD* | LPS Translocon Subunit D | Keio, PEC, TraDIS | presence gate | stops |
| *lptE* | LPS Translocon Subunit E | Keio, PEC, TraDIS | presence gate | stops |
| *lptG* | LPS Transport Permease | Keio, PEC, TraDIS | presence gate | stops |
| *mlaB* | Phospholipid Retrograde Transport (Mla System) | Keio | not wired | grows |
| *surA* | Periplasmic Chaperone (PPIase) | (none) | not wired | grows |
| *wzyE* | ECA Polymerase | Keio, TraDIS | presence gate | stops |
| **Peptidoglycan Biosynthesis** (16) | | | | |
| *ddlB* | D-Ala-D-Ala Ligase | (none) | not wired | grows |
| *mraY* | Phospho-MurNAc-Pentapeptide Translocase | Keio, PEC, TraDIS | presence gate | stops |
| *mrdA* | Penicillin-Binding Protein 2 (PBP2) | Keio, PEC, TraDIS | presence gate | stops |
| *mrdB* | Rod Shape-Determining Protein (RodA) | Keio, PEC, TraDIS | presence gate | stops |
| *mreB* | Actin Homolog / Rod-Shape Determinant | Keio, PEC, TraDIS | presence gate | stops |
| *mreC* | Elongasome Protein MreC | Keio, PEC, TraDIS | presence gate | – |
| *mreD* | Elongasome Protein MreD | Keio, PEC, TraDIS | presence gate | – |
| *murA* | UDP-GlcNAc Enolpyruvyl Transferase | Keio, PEC, TraDIS | presence gate | stops |
| *murB* | UDP-GlcNAc Enolpyruvyl Reductase | Keio, PEC, TraDIS | presence gate | stops |
| *murC* | UDP-MurNAc:L-Ala Ligase | Keio, PEC, TraDIS | presence gate | stops |
| *murD* | UDP-MurNAc-Dipeptide:D-Glu Ligase | Keio, PEC, TraDIS | presence gate | stops |
| *murE* | UDP-MurNAc-Tripeptide Ligase | Keio, PEC, TraDIS | presence gate | stops |
| *murF* | UDP-MurNAc-Pentapeptide Ligase | Keio, PEC, TraDIS | presence gate | stops |
| *murG* | GlcNAc Transferase | Keio, PEC, TraDIS | presence gate | stops |
| *murI* | Glutamate Racemase | Keio, PEC, TraDIS | presence gate | stops |
| *murJ* | Lipid II Flippase | Keio, PEC, TraDIS | presence gate | stops |
| **Phospholipid & Fatty Acid Biosynthesis** (21) | | | | |
| *accA* | Acetyl-CoA Carboxylase Subunit | Keio, PEC, TraDIS | presence gate | – |
| *accB* | Acetyl-CoA Carboxylase Biotin Carrier | Keio, PEC, TraDIS | presence gate | – |
| *accC* | Acetyl-CoA Carboxylase Biotin Carboxylase | Keio, PEC, TraDIS | presence gate | – |
| *accD* | Acetyl-CoA Carboxylase Subunit | Keio, PEC, TraDIS | presence gate | – |
| *acpP* | Acyl Carrier Protein | Keio, PEC, TraDIS | presence gate | – |
| *acpS* | Holo-ACP Synthase | Keio, PEC, TraDIS | presence gate | – |
| *cdsA* | CDP-Diacylglycerol Synthase | Keio, PEC, TraDIS | presence gate | – |
| *fabA* | Beta-Hydroxyacyl-ACP Dehydratase/Isomerase | Keio, PEC, TraDIS | presence gate | – |
| *fabB* | Beta-Ketoacyl-ACP Synthase I | Keio, PEC, TraDIS | presence gate | – |
| *fabD* | Malonyl-CoA:ACP Transacylase | Keio, PEC, TraDIS | presence gate | – |
| *fabG* | Beta-Ketoacyl-ACP Reductase | Keio, PEC, TraDIS | presence gate | – |
| *fabH* | Beta-Ketoacyl-ACP Synthase III | TraDIS | not wired | grows |
| *fabI* | Enoyl-ACP Reductase | Keio, PEC, TraDIS | presence gate | – |
| *fabZ* | Beta-Hydroxyacyl-ACP Dehydratase | Keio, PEC, TraDIS | presence gate | – |
| *gpsA* | Glycerol-3-Phosphate Dehydrogenase | Keio, PEC, TraDIS | reactions | grows |
| *pgsA* | Phosphatidylglycerophosphate Synthase | Keio, PEC, TraDIS | presence gate | – |
| *plsB* | Glycerol-3-Phosphate Acyltransferase | Keio, PEC, TraDIS | presence gate | – |
| *plsC* | 1-Acylglycerol-3-Phosphate Acyltransferase | Keio, PEC, TraDIS | presence gate | – |
| *plsX* | Acyl-ACP:Phosphate Acyltransferase | (none) | not wired | grows |
| *psd* | Phosphatidylserine Decarboxylase | Keio, PEC, TraDIS | presence gate | – |
| *pssA* | Phosphatidylserine Synthase | Keio, PEC, TraDIS | presence gate | – |
| **Protein Secretion & Membrane Insertion** (12) | | | | |
| *ffh* | Signal Recognition Particle Protein | Keio, PEC, TraDIS | presence gate | – |
| *ftsY* | SRP Receptor | Keio, PEC, TraDIS | presence gate | – |
| *lepB* | Signal Peptidase I | Keio, PEC, TraDIS | presence gate | – |
| *lgt* | Prolipoprotein Diacylglyceryl Transferase | Keio, PEC, TraDIS | presence gate | – |
| *lspA* | Lipoprotein Signal Peptidase | Keio, PEC, TraDIS | presence gate | – |
| *secA* | Protein Translocation ATPase | Keio, PEC, TraDIS | presence gate | – |
| *secD* | Translocation Accessory Factor | Keio, PEC | presence gate | – |
| *secE* | Translocon Subunit | Keio, PEC, TraDIS | presence gate | – |
| *secF* | Translocation Accessory Factor | Keio, PEC | presence gate | – |
| *secM* | Secretion Monitor | Keio, PEC | presence gate | – |
| *secY* | Protein-Conducting Channel | Keio, PEC, TraDIS | presence gate | – |
| *yidC* | Membrane Protein Insertase | Keio, PEC, TraDIS | presence gate | – |
| **RNA Processing & Degradation** (8) | | | | |
| *cca* | tRNA CCA-Adding Enzyme | Keio, PEC, TraDIS | presence gate | slows |
| *orn* | Oligoribonuclease | Keio, PEC, TraDIS | presence gate | – |
| *pnp* | Polynucleotide Phosphorylase | (none) | slowdown | – |
| *rnc* | RNase III | Keio | slowdown | grows |
| *rne* | RNase E | Keio, PEC | presence gate | slows |
| *rnpA* | RNase P Protein Subunit | Keio, PEC, TraDIS | presence gate | slows |
| *rpsB* | Ribosomal Protein S2 / mRNA turnover partner | Keio, PEC, TraDIS | subunit at assembly | – |
| *yceQ* | Small Reading Frame YceQ | Keio, PEC | presence gate | stops |
| **Respiratory Chain** (5) | | | | |
| *cydA* | Cytochrome bd-I Subunit CydA | Keio, TraDIS | presence gate | stops |
| *cydB* | Cytochrome bd-I Oxidase Subunit I | TraDIS | slowdown | slows |
| *cydC* | Cytochrome bd Assembly Transporter CydC | Keio, TraDIS | presence gate | – |
| *cydD* | Cytochrome bd Oxidase Assembly ABC Transporter | TraDIS | slowdown | – |
| *cydX* | Cytochrome bd-I Oxidase Accessory Subunit | TraDIS | slowdown | – |
| **Ribosome Assembly & Biogenesis GTPases** (13) | | | | |
| *der* | Ribosome Biogenesis GTPase (EngA) | Keio, PEC, TraDIS | presence gate | slows |
| *engB* | 50S Assembly GTPase EngB (YihA) | Keio, PEC, TraDIS | presence gate | slows |
| *era* | Ribosome Biogenesis GTPase | Keio, PEC, TraDIS | presence gate | slows |
| *nusB* | Antitermination Factor NusB | PEC, TraDIS | presence gate | slows |
| *obgE* | Ribosome Biogenesis GTPase (ObgE/CgtA) | Keio, PEC, TraDIS | presence gate | slows |
| *rbfA* | Ribosome-Binding Factor A | TraDIS | slowdown | grows |
| *rimM* | 16S rRNA Processing Factor | TraDIS | slowdown | – |
| *rimP* | Ribosome Maturation Factor | (none) | slowdown | – |
| *rluD* | Ribosomal Pseudouridine Synthase | TraDIS | slowdown | – |
| *rsgA* | Ribosome Biogenesis GTPase (RsgA) | PEC, TraDIS | slowdown | – |
| *suhB* | Antitermination Partner SuhB | Keio, PEC, TraDIS | presence gate | slows |
| *ybeY* | Ribosome Quality-Control Endonuclease | TraDIS | slowdown | – |
| *yqgF* | 16S rRNA Processing Factor | Keio, PEC, TraDIS | presence gate | slows |
| **Transcription** (14) | | | | |
| *cohE* | e14 Prophage Repressor | Keio, TraDIS | presence gate | – |
| *csrA* | Post-Transcriptional Regulator CsrA | Keio, PEC, TraDIS | presence gate | – |
| *dicA* | Qin Prophage Repressor | Keio, TraDIS | reactions | grows |
| *greA* | Transcript Cleavage Factor | (none) | subunit at assembly | – |
| *nusA* | Elongation/Termination Factor | Keio, PEC, TraDIS | subunit at assembly | – |
| *nusG* | Transcription-Translation Coupling Factor | Keio, PEC, TraDIS | subunit at assembly | – |
| *racR* | Rac Prophage Repressor | Keio, TraDIS | reactions | grows |
| *rho* | Transcription Termination Factor | PEC, TraDIS | subunit at assembly | – |
| *rpoA* | RNA Polymerase Alpha Subunit | Keio, PEC, TraDIS | subunit at assembly | – |
| *rpoB* | RNA Polymerase Beta Subunit | Keio, PEC, TraDIS | subunit at assembly | – |
| *rpoC* | RNA Polymerase Beta-Prime Subunit | Keio, PEC, TraDIS | subunit at assembly | – |
| *rpoD* | Sigma-70 (Housekeeping Sigma) | PEC, TraDIS | subunit at assembly | – |
| *rpoH* | Heat-Shock Sigma Factor | Keio, PEC, TraDIS | reactions | grows |
| *spoT* | ppGpp Synthase and Hydrolase | Keio, PEC | presence gate | stops |
| **Translation: 30S Ribosomal Proteins** (20) | | | | |
| *rpsA* | 30S Ribosomal Protein S1 | Keio, PEC, TraDIS | subunit at assembly | slows as recorded; ends at the wild-type count |
| *rpsC* | 30S Ribosomal Protein S3 | Keio, PEC, TraDIS | subunit at assembly | – |
| *rpsD* | 30S Ribosomal Protein S4 | Keio, PEC, TraDIS | subunit at assembly | – |
| *rpsE* | 30S Ribosomal Protein S5 | Keio, PEC, TraDIS | subunit at assembly | – |
| *rpsF* | 30S Ribosomal Protein S6 | TraDIS | subunit at assembly | – |
| *rpsG* | 30S Ribosomal Protein S7 | Keio, PEC, TraDIS | subunit at assembly | – |
| *rpsH* | 30S Ribosomal Protein S8 | Keio, PEC, TraDIS | subunit at assembly | – |
| *rpsI* | 30S Ribosomal Protein S9 | Keio, PEC, TraDIS | subunit at assembly | – |
| *rpsJ* | 30S Ribosomal Protein S10 | Keio, PEC, TraDIS | subunit at assembly | – |
| *rpsK* | 30S Ribosomal Protein S11 | Keio, PEC, TraDIS | subunit at assembly | – |
| *rpsL* | 30S Ribosomal Protein S12 | Keio, PEC, TraDIS | subunit at assembly | – |
| *rpsM* | 30S Ribosomal Protein S13 | Keio, PEC, TraDIS | subunit at assembly | – |
| *rpsN* | 30S Ribosomal Protein S14 | Keio, PEC, TraDIS | subunit at assembly | – |
| *rpsO* | 30S Ribosomal Protein S15 | TraDIS | subunit at assembly | – |
| *rpsP* | 30S Ribosomal Protein S16 | Keio, PEC, TraDIS | subunit at assembly | – |
| *rpsQ* | 30S Ribosomal Protein S17 | Keio, PEC, TraDIS | subunit at assembly | – |
| *rpsR* | 30S Ribosomal Protein S18 | Keio, PEC, TraDIS | subunit at assembly | – |
| *rpsS* | 30S Ribosomal Protein S19 | Keio, PEC, TraDIS | subunit at assembly | – |
| *rpsT* | 30S Ribosomal Protein S20 | TraDIS | subunit at assembly | – |
| *rpsU* | 30S Ribosomal Protein S21 | TraDIS | subunit at assembly | – |
| **Translation: 50S Ribosomal Proteins** (31) | | | | |
| *rplA* | 50S Ribosomal Protein L1 | TraDIS | subunit at assembly | grows |
| *rplB* | 50S Ribosomal Protein L2 | Keio, PEC, TraDIS | subunit at assembly | – |
| *rplC* | 50S Ribosomal Protein L3 | Keio, PEC, TraDIS | subunit at assembly | – |
| *rplD* | 50S Ribosomal Protein L4 | Keio, PEC, TraDIS | subunit at assembly | – |
| *rplE* | 50S Ribosomal Protein L5 | Keio, PEC, TraDIS | subunit at assembly | – |
| *rplF* | 50S Ribosomal Protein L6 | Keio, PEC, TraDIS | subunit at assembly | – |
| *rplI* | 50S Ribosomal Protein L9 | (none) | subunit at assembly | – |
| *rplJ* | 50S Ribosomal Protein L10 | Keio, PEC, TraDIS | subunit at assembly | – |
| *rplK* | 50S Ribosomal Protein L11 | TraDIS | subunit at assembly | – |
| *rplL* | 50S Ribosomal Protein L7/L12 | Keio, PEC, TraDIS | subunit at assembly | – |
| *rplM* | 50S Ribosomal Protein L13 | Keio, PEC, TraDIS | subunit at assembly | – |
| *rplN* | 50S Ribosomal Protein L14 | Keio, PEC, TraDIS | subunit at assembly | – |
| *rplO* | 50S Ribosomal Protein L15 | Keio, PEC, TraDIS | subunit at assembly | – |
| *rplP* | 50S Ribosomal Protein L16 | Keio, PEC, TraDIS | subunit at assembly | – |
| *rplQ* | 50S Ribosomal Protein L17 | Keio, PEC, TraDIS | subunit at assembly | – |
| *rplR* | 50S Ribosomal Protein L18 | Keio, PEC, TraDIS | subunit at assembly | – |
| *rplS* | 50S Ribosomal Protein L19 | Keio, PEC, TraDIS | subunit at assembly | – |
| *rplT* | 50S Ribosomal Protein L20 | Keio, PEC, TraDIS | subunit at assembly | – |
| *rplU* | 50S Ribosomal Protein L21 | Keio, PEC, TraDIS | subunit at assembly | – |
| *rplV* | 50S Ribosomal Protein L22 | Keio, PEC, TraDIS | subunit at assembly | – |
| *rplW* | 50S Ribosomal Protein L23 | Keio, PEC, TraDIS | subunit at assembly | – |
| *rplX* | 50S Ribosomal Protein L24 | Keio, PEC, TraDIS | subunit at assembly | – |
| *rplY* | 50S Ribosomal Protein L25 | TraDIS | subunit at assembly | – |
| *rpmA* | 50S Ribosomal Protein L27 | Keio, PEC, TraDIS | subunit at assembly | – |
| *rpmB* | 50S Ribosomal Protein L28 | Keio, PEC, TraDIS | subunit at assembly | – |
| *rpmC* | 50S Ribosomal Protein L29 | Keio, PEC, TraDIS | subunit at assembly | – |
| *rpmD* | 50S Ribosomal Protein L30 | Keio, PEC, TraDIS | subunit at assembly | – |
| *rpmE* | 50S Ribosomal Protein L31 | (none) | subunit at assembly | – |
| *rpmF* | 50S Ribosomal Protein L32 | TraDIS | subunit at assembly | – |
| *rpmH* | 50S Ribosomal Protein L34 | Keio, PEC, TraDIS | subunit at assembly | – |
| *rpmI* | 50S Ribosomal Protein L35 | TraDIS | subunit at assembly | – |
| **Translation: Factors & rRNA/tRNA** (18) | | | | |
| *def* | Peptide Deformylase | Keio, PEC, TraDIS | reactions | grows |
| *fmt* | Methionyl-tRNA Formyltransferase | Keio, PEC, TraDIS | presence gate | – |
| *frr* | Ribosome Recycling Factor | Keio, PEC, TraDIS | subunit at assembly | – |
| *fusA* | Elongation Factor G (EF-G) | Keio, PEC, TraDIS | subunit at assembly | – |
| *infA* | Initiation Factor IF1 | Keio, PEC, TraDIS | subunit at assembly | – |
| *infB* | Initiation Factor IF2 | Keio, PEC, TraDIS | subunit at assembly | – |
| *infC* | Initiation Factor IF3 | Keio, PEC, TraDIS | subunit at assembly | – |
| *map* | Methionine Aminopeptidase | Keio, PEC, TraDIS | reactions | grows |
| *pheM* | Phe Attenuator Leader Peptide | TraDIS | not wired | – |
| *prfA* | Release Factor 1 (RF1) | Keio, PEC, TraDIS | subunit at assembly | – |
| *prfB* | Release Factor 2 (RF2) | PEC, TraDIS | subunit at assembly | – |
| *prmC* | Release Factor Methyltransferase | Keio, PEC, TraDIS | presence gate | – |
| *pth* | Peptidyl-tRNA Hydrolase | Keio, PEC, TraDIS | presence gate | – |
| *rrfA* | 5S Ribosomal RNA | (none) | not wired | – |
| *rrlA* | 23S Ribosomal RNA | (none) | not wired | – |
| *rrsA* | 16S Ribosomal RNA | (none) | not wired | – |
| *tsf* | Elongation Factor Ts (EF-Ts) | Keio, PEC, TraDIS | subunit at assembly | – |
| *tufA* | Elongation Factor Tu (EF-Tu) | (none) | not wired | – |
| **tRNA Modification** (8) | | | | |
| *mnmA* | tRNA 2-Thiouridylase | TraDIS | slowdown | slows |
| *tadA* | tRNA-Specific Adenosine Deaminase | Keio, TraDIS | presence gate | stops |
| *tilS* | tRNA-Ile Lysidine Synthase | Keio, PEC, TraDIS | presence gate | stops |
| *trmD* | tRNA m1G37 Methyltransferase | Keio, PEC, TraDIS | presence gate | slows |
| *tsaB* | t6A Synthesis Subunit TsaB | Keio, PEC, TraDIS | presence gate | slows |
| *tsaC* | t6A Synthesis Enzyme TsaC | Keio, PEC, TraDIS | presence gate | slows |
| *tsaD* | t6A Synthesis Subunit TsaD | Keio, PEC, TraDIS | presence gate | slows |
| *tsaE* | t6A Synthesis Subunit TsaE | Keio, PEC, TraDIS | presence gate | slows |

#### Supplementary Table S2. The measured metabolite pools

Each cytoplasmic pool that Bennett *et al.* (2009) measured in absolute units in glucose-grown *E. coli*,
against the model averaged over seeds 1 and 2 from *t* = 1,000 to 3,000 s on *E. coli* version 1.0,
with the log_2_ fold change and the model species compared. Metabolites without a model species are marked.

| Metabolite | Measured | Model | log_2_(model/meas.) | Model species |
| --- | --- | --- | --- | --- |
| 2,3-dihydroxybenzoate | 140 µM | 41.7 µM | -1.75 | 2,3-dihydroxybenzoate |
| 3-phosphoglycerate | 1.5 mM | 756 µM | -0.99 | 3-phospho-D-glycerate |
| 4-hydroxybenzoate | 52 µM | 558 nM | -6.54 | 4-hydroxybenzoate |
| 6-phosphogluconate | 3.8 mM | 3.5 mM | -0.12 | 6-phospho-D-gluconate |
| acetoacetyl-CoA | 22 µM | 6.55 µM | -1.75 | acetoacetyl-CoA |
| acetyl-CoA | 610 µM | 425 µM | -0.52 | acetyl-CoA |
| acetylphosphate | 1.1 mM | 328 µM | -1.75 | acetyl-phosphate |
| adenine | 1.5 µM | 256 µM | +7.42 | adenine |
| adenosine | 130 nM | 216 µM | +10.70 | adenosine |
| adenosine-phosphosulfate | 6.6 µM | — | — | not in model |
| ADP | 560 µM | 579 µM | +0.05 | ADP |
| ADP-glucose | 4.3 µM | — | — | not in model |
| alanine | 2.6 mM | 3.15 mM | +0.28 | L-alanine |
| alpha-ketoglutarate | 440 µM | 953 µM | +1.11 | 2-oxoglutarate |
| AMP | 280 µM | 467 µM | +0.74 | AMP |
| anthranilate | 3.5 µM | 13.3 µM | +1.93 | anthranilate |
| arginine | 570 µM | 981 µM | +0.78 | L-arginine |
| asparagine | 510 µM | 2.13 mM | +2.06 | L-asparagine |
| aspartate | 4.2 mM | 8.03 mM | +0.94 | L-aspartate |
| ATP | 9.6 mM | 9.6 mM | +0.00 | ATP |
| carbamyl-aspartate | 590 µM | 25.7 µM | -4.52 | N-carbamoyl-L-aspartate |
| cis-aconitate | 16 µM | — | — | not in model |
| citrate | 2 mM | 362 µM | -2.47 | citrate |
| citrulline | 1.4 mM | 9.4 µM | -7.22 | L-citrulline |
| CMP | 360 µM | 544 µM | +0.60 | CMP |
| CoA | 1.4 mM | 1.4 mM | +0.00 | CoA |
| CTP | 2.7 mM | 4.06 mM | +0.59 | CTP |
| cyclic-AMP | 35 µM | 38.5 µM | +0.14 | cAMP |
| cytidine | 2.6 µM | 1.61 µM | -0.69 | cytidine |
| cytosine | 14 µM | 395 µM | +4.82 | cytosine |
| dAMP | 8.8 µM | 104 µM | +3.56 | dAMP |
| dATP | 16 µM | 169 µM | +3.40 | dATP |
| dCTP | 35 µM | 159 µM | +2.18 | dCTP |
| deoxyadenosine | 2.8 µM | 955 nM | -1.55 | deoxyadenosine |
| deoxyguanosine | 520 nM | 780 nM | +0.58 | deoxyguanosine |
| deoxyribose-5-P | 300 µM | 89.8 µM | -1.74 | 2-deoxy-D-ribose-5-phosphate |
| dGMP | 51 µM | 395 µM | +2.95 | dGMP |
| dihydroorotate | 12 µM | 12.4 µM | +0.05 | (S)-dihydroorotate |
| dihydroxyacetone-phosphate | 370 µM | 187 µM | -0.98 | dihydroxyacetone-phosphate |
| dTDP | 380 µM | 117 µM | -1.70 | dTDP |
| dTTP | 4.6 mM | 706 µM | -2.70 | dTTP |
| FAD | 170 µM | 627 µM | +1.88 | FAD |
| FMN | 54 µM | 427 nM | -6.98 | FMN |
| fructose-1,6-bisphosphate | 15 mM | 13.2 mM | -0.18 | D-fructose-1,6-bisphosphate |
| fumarate | 120 µM | 321 µM | +1.42 | fumarate |
| GDP | 680 µM | 1.74 mM | +1.36 | GDP |
| gluconate | 42 µM | 2.83 µM | -3.89 | D-gluconate |
| gluconolactone | 1 mM | 476 µM | -1.07 | 6-phospho-D-glucono-1,5-lactone |
| glucosamine-6-phosphate | 1.2 mM | — | — | not in model |
| glutamate | 96 mM | 10.6 mM | -3.18 | L-glutamate |
| glutamine | 3.8 mM | 4.82 mM | +0.34 | L-glutamine |
| glutathione | 17 mM | 10.9 mM | -0.64 | glutathione |
| glutathione-disulfide | 2.4 mM | — | — | not in model |
| glycerate | 1.4 mM | 416 µM | -1.75 | glycerate |
| glycerol-3-phosphate | 49 µM | 5.47 µM | -3.16 | glycerol-3-phosphate |
| GMP | 24 µM | 616 µM | +4.68 | GMP |
| GTP | 4.9 mM | 6.17 mM | +0.33 | GTP |
| guanine | 190 µM | 31.4 µM | -2.60 | guanine |
| guanosine | 1.6 µM | 2.43 µM | +0.60 | guanosine |
| hexose-P | 8.8 mM | 3.89 mM | -1.18 | D-glucose-6-phosphate+D-fructose-6-phosphate |
| histidine | 68 µM | 95.2 µM | +0.49 | L-histidine |
| histidinol | 13 µM | 57 µM | +2.13 | L-histidinol |
| homocysteine | 370 µM | 72.5 µM | -2.35 | L-homocysteine |
| IMP | 270 µM | 482 µM | +0.84 | IMP |
| inosine-diphosphate | 24 µM | 68.5 µM | +1.51 | IDP |
| inosine-triphosphate | 210 µM | 60.8 µM | -1.79 | ITP |
| isoleucine+leucine | 300 µM | 572 µM | +0.93 | L-isoleucine+L-leucine |
| lysine | 410 µM | 710 µM | +0.79 | L-lysine |
| malate | 1.7 mM | 3.18 mM | +0.90 | L-malate |
| malonyl-CoA | 35 µM | 10.2 µM | -1.78 | malonyl-CoA |
| methionine | 150 µM | 104 µM | -0.53 | L-methionine |
| myo-inositol | 5.7 µM | 1.7 µM | -1.75 | myo-inositol |
| N-acetyl-glucosamine-1P | 82 µM | — | — | not in model |
| N-acetyl-ornithine | 43 µM | 1.14 µM | -5.24 | N2-acetyl-L-ornithine |
| NAD+ | 2.6 mM | 3.03 mM | +0.22 | NAD+ |
| NADH | 83 µM | 438 µM | +2.40 | NADH |
| NADP+ | 2.1 µM | 1.3 mM | +9.27 | NADP+ |
| NADPH | 120 µM | 213 µM | +0.83 | NADPH |
| ornithine | 10 µM | 2.49 µM | -2.01 | L-ornithine |
| pentose-P | 1.3 mM | 616 µM | -1.08 | D-ribulose-5-phosphate+alpha-D-ribose-5-phosphate+D-xylulose-5-phosphate |
| phenylalanine | 18 µM | 39.1 µM | +1.12 | L-phenylalanine |
| phenylpyruvate | 90 µM | 30.3 µM | -1.57 | phenylpyruvate |
| phosphoenolpyruvate | 180 µM | 229 µM | +0.35 | phosphoenolpyruvate |
| proline | 390 µM | 551 µM | +0.50 | L-proline |
| propionyl-CoA | 5.3 µM | 1.58 µM | -1.75 | propionyl-CoA |
| PRPP | 260 µM | 228 µM | -0.19 | 5-phospho-alpha-D-ribose-1-diphosphate |
| quinolinate | 12 µM | 119 µM | +3.31 | quinolinate |
| riboflavin | 19 µM | 4.93 µM | -1.95 | riboflavin |
| S-adenosyl-L-methionine | 180 µM | 180 µM | +0.00 | S-adenosyl-L-methionine |
| serine | 68 µM | 58.8 µM | -0.21 | L-serine |
| shikimate | 14 µM | 140 µM | +3.32 | shikimate |
| succinate | 570 µM | 1.31 mM | +1.20 | succinate |
| succinyl-CoA | 230 µM | 453 µM | +0.98 | succinyl-CoA |
| threonine | 180 µM | 128 µM | -0.49 | L-threonine |
| tryptophan | 12 µM | 35 µM | +1.54 | L-tryptophan |
| tyrosine | 29 µM | 56 µM | +0.95 | L-tyrosine |
| UDP | 1.8 mM | 2.73 mM | +0.60 | UDP |
| UDP-glucose | 2.5 mM | — | — | not in model |
| UDP-glucuronate | 570 µM | — | — | not in model |
| UDP-N-acetyl-glucosamine | 9.2 mM | 8.58 mM | -0.10 | UDP-N-acetyl-D-glucosamine |
| uridine | 2.1 mM | 625 µM | -1.75 | uridine |
| UTP | 8.3 mM | 12.5 mM | +0.59 | UTP |
| valine | 4 mM | 767 µM | -2.38 | L-valine |

#### Supplementary Table S3. The 31 pools held at fixed values

The 31 pools held at fixed values in *E. coli* version 1.0, with the values as compiled. Ten are held by
design: H^+^, O_2_ and CO_2_ in the medium, the periplasm and the cytoplasm, which supply
protons and the two gases at a constant level (the H^+^ value is a supply pool and does not represent pH), and
the loaded replication origin, since initiation of replication is not modelled. Phosphate is held at its measured
cytoplasmic level. The other 20 stand for synthesis not yet written; three of them (carbamoyl phosphate and the two
ThiS carrier forms) were released after version 1.0 as their pathways were written and tested in the whole cell.

| Pool | Compartment | Held value | Status |
| --- | --- | --- | --- |
| H^+^ | medium | 1.2 mM | by design: proton supply |
| O_2_ | medium | 1.2 mM | by design: gas supply |
| CO_2_ | medium | 1.2 mM | by design: gas supply |
| H^+^ | periplasm | 1.2 mM | by design: proton supply |
| O_2_ | periplasm | 1.2 mM | by design: gas supply |
| CO_2_ | periplasm | 1.2 mM | by design: gas supply |
| H^+^ | cytoplasm | 1.2 mM | by design: proton supply (with a ProtonSupply row) |
| O_2_ | cytoplasm | 1.2 mM | by design: gas supply |
| CO_2_ | cytoplasm | 1.2 mM | by design: gas supply |
| oriC~DNAP (loaded origin) | cytoplasm | 1 | by design: initiation not modelled |
| phosphate | cytoplasm | 23.9 mM | held at its measured level; no reason recorded |
| coenzyme A | cytoplasm | 1.4 mM | waits for CoA synthesis under feedback |
| bicarbonate | cytoplasm | 2 mM | waits for its own supply and use |
| ammonium | cytoplasm | 2 mM | waits for ammonium uptake and assimilation balance |
| ribose 5-phosphate | cytoplasm | 0.4 mM | waits for the pentose phosphate pathway |
| ribulose 5-phosphate | cytoplasm | 0.16 mM | waits for the pentose phosphate pathway |
| erythrose 4-phosphate | cytoplasm | 97 µM | waits for the pentose phosphate pathway |
| pyridoxal 5'-phosphate | cytoplasm | 100 µM | waits for vitamin B6 synthesis |
| pyridoxamine 5'-phosphate | cytoplasm | 100 µM | waits for vitamin B6 synthesis |
| S-adenosylmethionine | cytoplasm | 180 µM | waits for SAM synthesis and use |
| tetrahydrofolate | cytoplasm | 200 µM | waits for purine control (released once, then held again) |
| 5,10-methylenetetrahydrofolate | cytoplasm | 200 µM | as tetrahydrofolate |
| 7,8-dihydrofolate | cytoplasm | 200 µM | as tetrahydrofolate |
| 10-formyltetrahydrofolate | cytoplasm | 200 µM | as tetrahydrofolate |
| 5-methyltetrahydrofolate | cytoplasm | 200 µM | as tetrahydrofolate |
| meso-diaminopimelate | cytoplasm | 80 µM | waits for diaminopimelate synthesis in the cell wall unit |
| farnesyl diphosphate | cytoplasm | 0.4 µM | waits for isoprenoid synthesis |
| isopentenyl diphosphate | cytoplasm | 4.1 µM | waits for isoprenoid synthesis |
| carbamoyl phosphate | cytoplasm | 200 µM | released after version 1.0 |
| ThiS-Gly-Gly (sulfur carrier) | cytoplasm | 100 µM | released after version 1.0 |
| ThiS thiocarboxylate | cytoplasm | 100 µM | released after version 1.0 |
